# Large-scale identification of RBP-RNA interactions by RAPseq refines essentials of post-transcriptional gene regulation

**DOI:** 10.1101/2021.11.08.467743

**Authors:** Ionut Atanasoai, Sofia Papavasileiou, Natalie Preiß, Claudia Kutter

**Affiliations:** Department of Microbiology, Tumor and Cell Biology, Karolinska Institute, Science for Life Laboratory, Sweden

**Keywords:** RNA binding proteins, RBP cooperativity, non-canonical RBPs, RBP mutants, RBP isoforms, native transcriptome, RNA modifications, RNA sequencing, RNA processing, vertebrate evolution

## Abstract

Over the past decade, thousands of putative human RNA binding proteins (RBPs) have been identified and increased the demand for specifying RNA binding capacities. Here, we developed RNA affinity purification followed by sequencing (RAPseq) that enables *in vitro* large-scale profiling of RBP binding to native RNAs. First, by employing RAPseq, we found that vertebrate HURs recognize a conserved RNA binding motif and bind predominantly to introns in zebrafish compared to 3’UTRs in human RNAs. Second, our dual RBP assays (co-RAPseq) uncovered cooperative RNA binding of HUR and PTBP1 within an optimal distance of 27 nucleotides. Third, we developed T7-RAPseq to discern m^6^A-dependent and - independent RNA binding sites of YTHDF1. Fourth, RAPseq of 26 novel non-canonical RBPs revealed specialized moonlighting interactions. Last, five pathological IGF2BP family variants exhibited different RNA binding patterns. Overall, our simple, scalable and versatile method enables to fast-forward RBP-related questions.

**Graphical Abstract:** 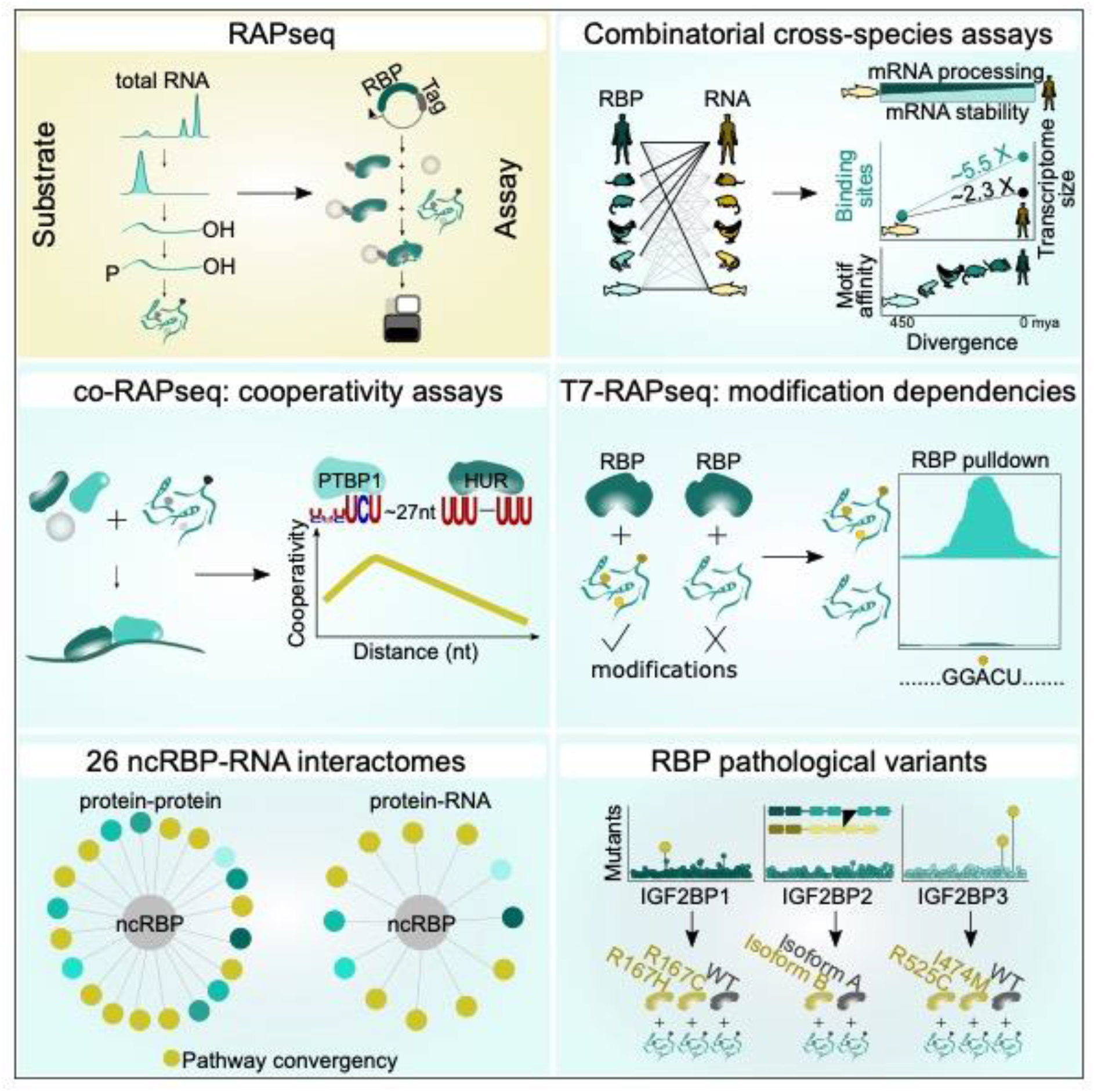

**HIGHLIGHTS:**

- RAPseq reveals *in vitro*-derived RBP-RNA interactomes
- the vertebrate-conserved HUR binding motif adapted to species-unique RNA features
- co-RAPseq and T7-RAPseq uncover binding cooperativity and modification dependencies
- non-canonical RBPs have specialized RNA interactomes

## INTRODUCTION

RNA binding proteins (RBPs) decide the fate and function of all RNA species from synthesis to decay. Latest advances of crosslinking-based methods coupled with mass-spectrometry enabled the detection of RBP-RNA complexes and have extended the number of putative human RBPs to about 5,000 (Baltz et al., 2012; Castello et al., 2012, 2016; Queiroz et al., 2019; Trendel et al., 2019; Urdaneta et al., 2019). These can be grouped into canonical (c) or non-canonical (nc) RBPs depending on the presence or absence of a known RNA binding domain, respectively. Furthermore, technological developments allowing transcriptome-wide mapping of modified cellular RNA species uncovered extensive changes that impact binding of RBPs (Alarcón et al., 2015; Dominissini et al., 2012; Dominissini et al., 2016; Huang et al., 2018; Meyer, 2019; Wu et al., 2018; L.-S. Zhang et al., 2019).

To uncover mechanisms of RBP-mediated regulatory processes, it is crucial to identify RBP-bound RNA molecules, their exact RNA binding sites and interaction strengths. UV-crosslinking followed by immunoprecipitation (CLIP) (Ule et al., 2003) has revolutionized detecting RBP binding sites, and by coupling it to high-throughput RNA sequencing (RNAseq) has yielded many variations of the method, including HITS-CLIP (Licatalosi et al., 2008), PAR-CLIP (Hafner et al., 2010), iCLIP (König et al., 2010), irCLIP (Zarnegar et al., 2016) and eCLIP (Van Nostrand et al., 2016), among others. Although CLIPseq methods have enabled breakthroughs in the field, limitations remain due to the low UV-crosslinking efficiency favoring distinct amino acid-ribonucleotide interactions, antibody availability and extensive hands-on time to produce CLIPseq libraries resulting in quantitative errors and high experimental failure rates (Lin & Miles, 2019; Wheeler et al., 2018). In addition, large-scale *in vitro* technologies, such as RNA Bind-n-Seq (RBNS) (Lambert et al., 2014), RNAcompete (Ray et al., 2013) and HTR-SELEX (Jolma et al., 2020), were developed to profile binding of purified recombinant RBPs to random synthetic RNAs of 20 or 40 nucleotides (nt) and allow quantitative assessment of primary and secondary binding motifs. However, these methods cannot infer the impact of intrinsic RNA features present in native cellular transcripts on RNA binding affinities.

To enable quantitative assessments of RBP binding sites in cellular transcriptomes, we developed RAPseq, an RBP-centric affinity purification-based method. RAPseq allows to study *in vitro* RBP-RNA interactomes. We applied RAPseq to (i) describe the evolution of HUR as an essential RBP across multiple vertebrates by cross-species RAPseq assays, (ii) measure transcriptome-wide cooperativity of HUR and PTBP1 through co-RAPseq assays, (iii) clarify the role of YTHDF1 as an m^6^A-reader by comparing cellular (RAPseq) and *in vitro* generated (T7-RAPseq) transcriptomes, (iv) profile transcriptomes bound by 26 ncRBPs and (v) assess the impact of disease-associated RBP variants in RNA binding activities on a transcriptome-wide scale. Altogether, RAPseq proves to be a simple, scalable and multiplexable method that can be easily implemented to deliver robust and highly reproducible RBP-RNA interactomes across various applications.

## RESULTS

### RAPseq enables interrogating RBP-RNA interactions based on RNA structure-, sequence- and modification-specific binding modalities

RAPseq is a binding assay that measures interactions between a recombinant protein and native RNA using next generation sequencing (NGS) as a readout (**Figure 1A**). For generating the RNA substrate, we chemically fragmented RNA isolated from cells and tissues to circa 35 nt products (**Figure S1A**). The subsequent RNA 3’-end dephosphorylation and 5’-end phosphorylation enabled ligation of NGS compatible adapters (**Figure 1A**). For producing the recombinant RBP, we cloned the respective cDNA fused to HaloTag (Los et al., 2008) into a cell-free expression vector (**Figure 1A**). Tagging Halo at the RBP C-terminus ensured assaying full-length rather than N-terminally tagged truncated fusions (**Figure S1B-C)**. We RNase A-treated the RBP from the *in vitro* expression system to eliminate residual RNA (**Figure S1D**) and removed contaminants by high-salt washes. After purification, the RBP-Halo fusion was incubated with the fragmented RNA substrate, the bound molecules were recovered and deep sequenced (**Figure 1A**) (STAR Methods). We also tested full-length total RNA as substrate for the binding assay. Unlike full-length, the fragmented RNA substrate improved the RBP-RNA pulldown efficiency (**Figure 1B**).

**Figure 1.**
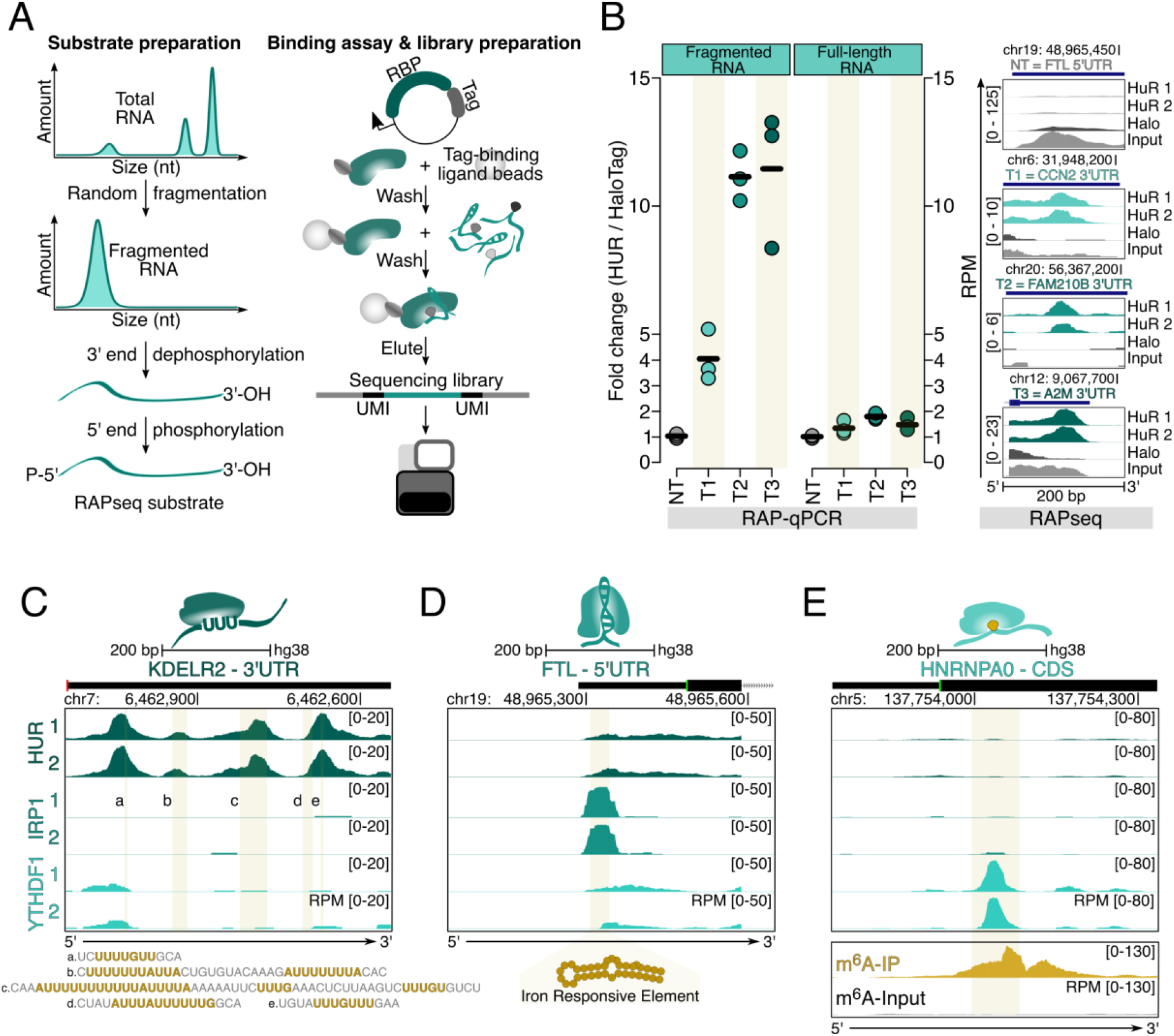
RAPseq profiles RBPs recognizing RNA structure, sequence, and modification. **(A)** Schematic presentation of the experimental setup of RAPseq. RAPseq substrates are generated from native total RNA (left). Production of a RBP-Halo fusion allows protein purification and RNA binding assay. Next-generation sequencing serves as read-out using unique molecular identifiers (UMI) (right). **(B)** Dotplot displays binding of HUR to non-targeting (NT) and targeting (T1 to T3) RNA regions (x-axis) of fragmented (left) and full-length (right) RNA from HepG2 cells. RAP-qPCR (left) was performed in three replicates (colored dots), means (black horizontal lines) are shown. RNA binding is displayed as fold change (ΔΔCt) of HUR over HaloTag control (y-axis). RAPseq coverage tracks (right) represent HUR binding to fragmented RNA (in reads per million, RPM). Genomic locations of the corresponding target regions (T1-3) are specified. Tracks for two HUR replicates (green) and two RAPseq controls (HaloTag and RNA input, grey) are shown. (**C-E**) Genome tracks demonstrate binding of (**C**) HUR to ARE and GRE motifs, (**D**) IRP1 to the IRE and (**E**) YTHDF1 to a modified nucleotide (m^6^A) within KDELR2, FTL and HNRNPA0 mRNAs, respectively, in HepG2 cells. Genomic locations with scale bar indicate the length of the genomic region in bases and gene features (black rectangle: exon and UTR, grey line: intron, arrow: direction of transcription) (x-axis) and normalized read density (RPM, y-axis) are shown. Vertical yellow lines highlight bound RNA elements. Tracks for two RBP replicates (green) and m^6^A-specific RNA immunoprecipitation (IP, yellow) over input control are presented.

To assess whether RAPseq captured known RBP binding modalities to RNA, we profiled three RBPs that have sequence-(HUR) (Lebedeva et al., 2011; Ray et al., 2009; Ripin et al., 2019; Wang et al., 2013), structure-(IRP1) (Hentze et al., 1987) and modification-specific (YTHDF1) (D. Han et al., 2019; X. Wang et al., 2015; Xu et al., 2014) substrate requirements. In our RAPseq assays, we detected a cluster of HUR bound AU- and GU-rich RNA elements (ARE, GRE) bound by HUR in the 3’UTR of KDELR2 mRNA (**Figure 1C**). IRP1 bound the iron-responsive element (IRE) in the 5’UTR of FTL (**Figure 1D**). The YTHDF1 RNA binding site overlapped with a previously identified m^6^A site (Dominissini et al., 2012) (**Figure 1E**). In sum, RAPseq is the first *in vitro*-based method that captures all major forms of interactions between RBP and native cellular RNA.

### RAPseq confirms known RBP binding specificities to RNA

To benchmark RAPseq, we focused on five cRBPs (HNRNPA1, HNRNPC, PTBP1, RBFOX2 and YBX3) that recognize RNA through sequence-specific interactions. We profiled their RBP-RNA interactomes using HepG2 RNA and compared our results to publicly available *in cellulo* eCLIPseq data (Van Nostrand et al., 2020) and *in vitro* RBP-RNA interaction assays (RNA compete, RBNS and HTR-SELEX).

We observed high similarity in the bound motifs for the five RBPs across the diverse interaction assays (**Figure 2A** and **Figure S2A**). To assess the quantitative nature of RAPseq, we calculated a peak binding score, which is defined as the average enrichment of reads over our two negative controls weighted by the average replicate normalized read counts (STAR Methods). When determining the peak binding score as a function of uracil content at each individual HNRNPC binding site, our RAPseq data showed that binding of HNRNPC to its RNA targets reached a plateau at an uracil content of ≥50% per target. This trend was also visible when considering only fold changes demonstrating that RAPseq can produce typical biochemical binding curves. In contrast, this assessment was not detectable in eCLIP data (**Figure 2B**). For RBFOX2, we noticed a slight deviation from the GCAUG motif identified in the other assays. We found that the second position in the motif showed an equal abundance of C and A. This GAAUG motif has been recently reported (Begg et al., 2020). A more generalizable RBFOX2 motif, GNWYG (N corresponds to any, W to A/U, Y to C/U nucleotide), identified using lenient criteria (STAR Methods) was present in almost 90% of all binding sites compared to 44% for G(C/A)AUG. We observed that binding sites containing only the canonical GCAUG motif have significantly higher peak binding scores compared to the other 13 possible 5-mers confirming GCAUG as the motif with the highest binding affinity (**Figure S2B**). All GNWYG 5-mers were specific for RBFOX2 compared to our other RAPseq assays (**Figure S2C**). Peak binding scores for the majority (86%, 12/14) of the individual RBFOX2 5-mer motif increased upon consideration of an additional 5’U (**Figure S2D**) verifying the previously reported increase in binding affinity (Lambert et al., 2014). Binding sites containing two 5-mers had higher peak binding scores than sites with only one 5-mer (**Figure S2D**). We also confirmed the generalizable GNWYG motif when inspecting eCLIP enrichments as a function of the number of GNWYG (excluding GCAUG) motifs per peak (**Figure S2E**). Like RBFOX2, YBX3 peak binding scores increased in the presence of additional 5’ adenosines with respect to the core CAHC motif (**Figure S2F**).

**Figure 2.**
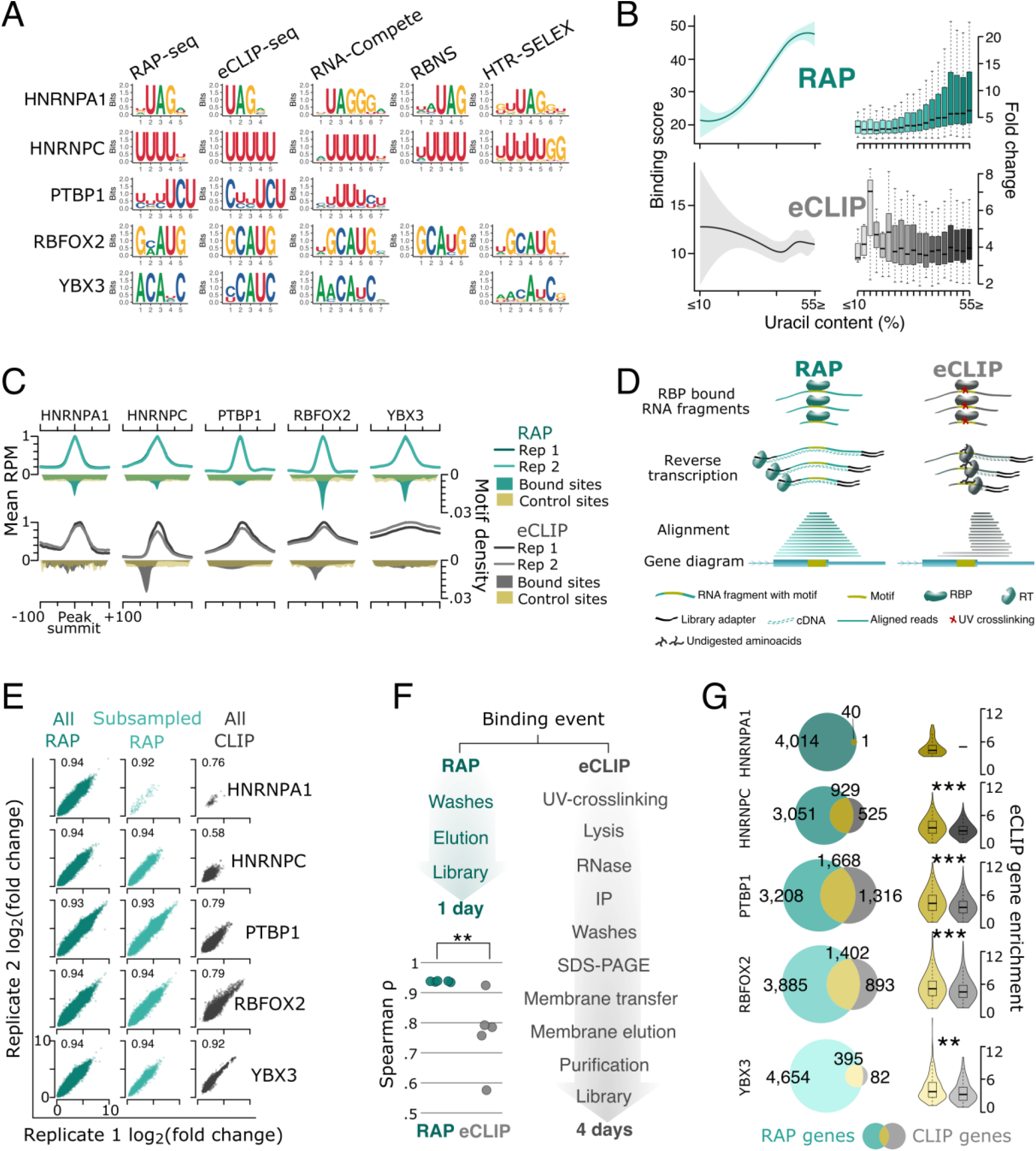
The robustness of RAPseq is confirmed by benchmarking against state-of-the-art methods. **(A)** Position weight matrices display *de novo* identified motifs of five RBPs in RAPseq and eCLIPseq assays and published motif models from three *in vitro* assays (RNAcompete, RBNS and HTR-SELEX). **(B)** LOESS fitted smoothened regression lines (left) and boxplots (right) of HNRNPC RAPseq (green, top) and eCLIPseq (grey, bottom) assays show HNRNPC binding in terms of peak binding scores (left) and fold changes (right) (y-axes) as a function of uracil content (x-axes) in a 40 nt bin centered around the peak summit (for RAPseq, top) and around the highest motif density location (for eCLIPseq, bottom). **(C)** Line and polygon peak metaplots compare the motif locations (polygons) with respect to the average normalized read enrichment (RPM, lines) for RAPseq (green) and eCLIPseq (grey) for five RBPs. Line plots show the average RPM for all peaks plotted as a fraction of the maximum mean RPM for two experimental replicates (top, y-axes). The bin size for the line plots is 1 nt. Polygon plots display the density of RBP binding motif locations (Figure 2A) at bound sites for RAPseq (green), eCLIPseq (grey) and the same motifs over unbound control sites (yellow). The x-axes show the distance from the peak summit. Ticks are drawn at -100, -50, 0, 50 and 100 nt from the respective RBP peak summit. **(D)** Illustrative model explains the read enrichment at (RAPseq, left) and next to the motif (eCLIPseq, right) due to intrinsic experimental differences when capturing RBP-RNA interactions. **(E)** Scatter plots show the correlations in binding enrichments (log_2_ of peak fold changes) between two replicates (x, y-axes) for all RAPseq binding events (left column) and subsampled RAPseq binding events (middle column) to match the number of eCLIP binding events (right column) for five RBPs (Spearman’s rank correlation coefficients). **(F)** Comparison of the experimental steps in RAP (left) and eCLIP protocols. Dotplot correlates the mean of two replicates for five RAPseq (green) and eCLIPseq (grey) experiments. Asterisks indicate *p*-values (Spearman rank correlation, **p<0.01). **(G)** Two-way Venn diagrams (left) intersect the number of genes with identified binding sites for five RBPs in RAPseq (green) and eCLIPseq (grey). The intersected area is colored (yellow) and the number of genes per intersection is displayed. Violin plots (right) of eCLIPseq demonstrate gene enrichments (in log_2_ scale) common in RAPseq and eCLIPseq (yellow) and eCLIPseq only genes (grey). Asterisks represent *p*-values (one-tailed Wilcoxon rank sum test, **p<0.01, ***p<0.001). The gene enrichments are computed as the sum of all peak enrichments (binding score) per gene.

We next compared the concordance between RAPseq and eCLIPseq regarding motif location and read enrichment of the five RBPs. In RAPseq, the motif locations were consistent with read densities around the center of peak coordinates resembling profiles from transcription factor chromatin immunoprecipitation (ChIP) assays. This allowed the use of standardized peak calling algorisms (STAR Methods). In contrast, CLIPseq methods require assay-dependent customized algorithms (Hafner et al., 2021) due to the shift between read and motif density locations (**Figure 2C-D**). RAPseq exhibited higher reproducibility and more significant binding events than eCLIPseq (Spearman rank correlation for RAPseq ρ=0.93-0.94 and for eCLIPseq ρ=0.58-0.92, Wilcoxon rank sum test for eCLIPseq and RAPseq p<0.01) (**Figure 2E-F)**. When combining peaks on the gene level, we noticed that the majority (56% to 83%) of RBP bound genes in eCLIPseq overlapped with RAPseq. More genes were bound by RBPs in our RAPseq assays when compared to eCLIPseq. However, binding of most of the transcripts *in cellulo* (eCLIPseq) can be attributed solely to the native RBP binding properties (RAPseq) (**Figure 2G**). In addition, transcripts bound *in vitro* were significantly stronger bound *in cellulo* (Wilcoxon rank sum test, p<0.01) (**Figure 2G**). Overall, RAPseq profiles quantitatively RBP-binding specificities and RBP-RNA interactomes.

### HUR binding to uracil triplets is conserved but has species-specific transcript-processing preferences

Species radiation underlines rapid changes in the regulation of the transcriptome requiring adaptation of RBP-RNA binding. RAPseq enables cross-species comparisons between RBP orthologs and RNA substrates. Since HUR is the most divergent paralog of the ELAV gene family that evolved through gene duplication in vertebrates (Good, 1995), we quantified and compared HUR binding specificities in vertebrate evolution. We inspected HUR orthologs from human, mouse, opossum, chicken, frog and fish. All orthologs had a similar length and the amino acid sequence identity changed gradually from human to zebrafish (**Figure 3A**). Except for the frog ortholog, all amino acid changes resided outside the RNA interacting beta sheets of the three RNA recognition motifs (RRMs) suggesting no or only few differences in RNA binding (**Figure S3A**). Despite two amino acid differences in the RRM1 and RRM3 beta sheets of the frog HUR, RNA contacts should be maintained according to crystallographic models showing specific interactions of all three RRMs with uracil triplets to the human structure (Ripin et al., 2019; H. Wang et al., 2013) (**Figure S3A**). To validate that binding specificity is conserved, we first produced HUR proteins of the tetrapod orthologs and performed RAPseq assays with the same human RNA substrate (**Figure 3B**). We found that tetrapod HURs bound to uracil triplets (UUU), which were significantly enriched in more than 90% of all ortholog binding events (**Figure 3C** and **S3A**). The uracil triplet was localized centrally at the peak summit over background control sites (**Figure 3D)** (STAR Methods). The fold changes were proportional to the number of triplets per binding site implying tetrapod-conserved RRM cooperativity in substrate recognition (**Figure 3E)**. However, we noticed differences in global enrichment profiles. The frog HUR had the lowest enrichments compared to the other tetrapod HURs suggesting that some of the amino acid changes caused differences in RNA binding affinities (Wilcoxon rank sum test, p<0.001) (**Figure 3F)**.

**Figure 3.**
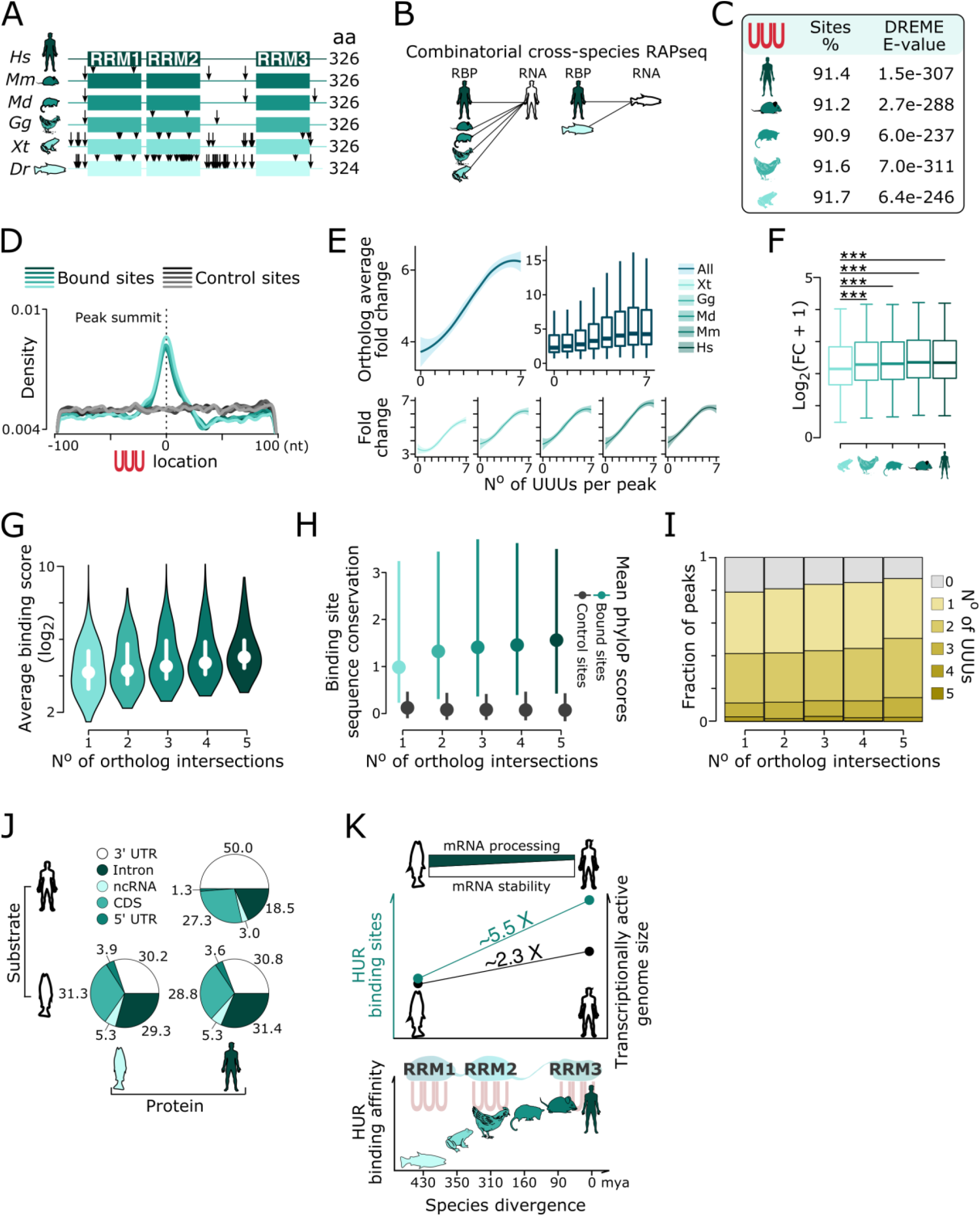
RNA binding specificity of HUR is conserved in vertebrates but acquired functional differences. **(A)** Orthologous vertebrate HUR protein length and RNA recognition motifs (RRM1-3) (green boxes) are shown (color-coded by species). Amino acid (aa) changes (black arrows) for each ortholog compared to the human reference are highlighted. **(B)** Schematic representation of the performed RAPseq assays using various vertebrate HUR proteins (colored) and either the human (left) or zebrafish RNA substrate (right). **(C)** Table lists the percentage and corresponding DREME E-value of uracil triplet motifs in the HUR-bound transcriptome (±25 nt around the peak summit). **(D)** Line plot (smoothing band width 2 nt) shows frequency of uracil triplets ± 100 nt at HURs peak summits (green, color-code by species) and control sites residing 1000 nt distant from the peak summit (grey, color-code by species). **(E)** Binding site enrichments (peak fold change, y-axis) are shown as a function of the frequency of uracil triplets (x-axis) for all HUR orthologs. LOESS fitted smoothened regression lines with 95% confidence interval display the averaged enrichments (Figure S3C) across all (top left) and for each individual ortholog (bottom panels). Boxplots (top right) show enrichment distributions (orthologue average fold change) of each uracil triplet bin. **(F)** Boxplots show peak fold change per ortholog (color-coded by species). Asterisks represent *p*-values (one-tailed Wilcoxon rank sum test, ***p<0.001). (**G-I**) Plots represent (**G**) average HUR binding (scored as log_2_ transformed average binding score), (**H**) degree of sequence conservation of HUR bound sites compared to adjacent unbound control sites (according to 100 vertebrate phyloP scores) and (**I**) proportional frequency (0-5) of uracil triplet motifs within the bound sites per orthologous species intersections (1-5) (Figure S3B). **(J)** Pie chart shows the percentage of HUR binding sites for a given RNA feature (color-coded) in the combinatorial human and zebrafish cross-species RAPseq assays. **(K)** Proposed model summarizes the co-evolutionary dependencies between HUR protein and transcriptomes in vertebrates. Top: From zebrafish to human, HUR binding transitioned from mRNA processing to mRNA stability (inverse triangles). Middle: During evolution, the size of the actively transcribed genome doubled from zebrafish to human while HUR binding sites increased 5-fold (line plots). Bottom: HUR binding specificity for uracil triplets remained unchanged (motif in the background) but HUR binding affinities changed across vertebrates (colored species cartoon). Hs, human; Mm, mouse; Md, opossum; Gg, chicken; Xt, frog; and Dr, zebrafish.

We grouped HUR binding by orthologous species intersections (**Figure S3B**) and averaged the peak binding scores of the tetrapod HURs for all RNA binding events to quantify the extent of binding similarities across all orthologs (**Figure S3C**). Average peak binding scores were higher in the tetrapod-conserved than species-specific RNA binding events (**Figure 3G)**. To assess the degree of evolutionary conservation of HUR binding sites, we used phyloP scores obtained from 100 vertebrate genome alignments. We found that ortholog-shared and - unique RNA binding sites were highly constrained compared to adjacent RNA sequences used as control (**Figure 3H**) (STAR Methods). Remarkably, the underlying sequence was under stronger selection in the common than the ortholog-unique RNA binding sites (**Figure 3H**) and followed the same trend as the average peak binding scores (**Figure 3G)** and uracil triplet count (**Figure 3I**). This suggested positive selection of uracil triplet expansion serving as HUR binding sites in the vertebrate lineage.

Given that the zebrafish HUR is most diverse across the vertebrate lineage, we profiled zebrafish and human HUR proteins using zebrafish RNA as substrate (**Figure 3B**). We found that the zebrafish and human HURs recognized uracil triplets in the zebrafish RNA with similar specificity and cooperativity (**Figure S3D-F)**. However, the zebrafish HUR had significantly lower enrichments than the human HUR (**Figure S3G**). By inspecting HUR binding events across the transcript body, we noticed a higher fraction of intronic sites in the zebrafish transcriptome compared to human, where most HUR sites reside in 3’UTRs (**Figure 3J**). To determine pathways that could be affected by this divergence, we examined the 94 Gene Ontology (GO) terms commonly bound in human and zebrafish transcriptomes. In this common set, the binding of zebrafish HUR occurred at a nearly even proportional frequency to zebrafish intronic (47%) and 3’UTR (53%) sequences whereas human HUR binding to human substrate was more prevalent in human 3’ UTRs (75%) relative to introns (25%) (**Figure S3H**). We next inspected the proportion of HUR binding sites within introns and 3’UTRs for each of the 94 GO terms (**Figure S3I**) and discovered the highest divergence in gene products mediating cellular responses to insulin in which 80% of HUR binding sites occurred in introns in zebrafish compared to 20% in humans (**Figure S3J**). Given that HUR is the only paralog with both nuclear and cytoplasmic functions in vertebrates and the original strictly nuclear function of ELAV proteins in invertebrates (Good, 1995), our results suggest an evolutionary shift in HUR function.

In conclusion, our cross-species combinatorial RAPseq experiments revealed vertebrate-conserved HUR binding to uracil triplets. Vertebrate HUR binding sites expanded selectively and affinities gradually increased from zebrafish to human. This allowed HUR to adapt predominant post-transcriptional roles from mRNA processing in zebrafish to mRNA stability in human (**Figure 3K**).

### HUR and PTBP1 cooperate post-transcriptionally by keeping a specific distance between binding sites

Previous reports have shown that HUR and PTBP1 act cooperatively and competitively to regulate mRNA processing (Galbán et al., 2008; Shwetha et al., 2015). In our assays, HUR and PTBP1 sites overlapped with a bimodal distance distribution separated at 30 nt. Proximal sites (0-30 nt, n=797) could result in competitive binding events, whereas distal peaks (31-100 nt, n=422) indicated co-binding (**Figure 4A**). We statistically inferred the outcome of competitive events and quantified that 17% and 22% of the proximal peaks could be significantly attributed to one or the other RBP (fold change ≥2, FDR-adjusted *p*≤0.05) (**Figure 4B**).

**Figure 4.**
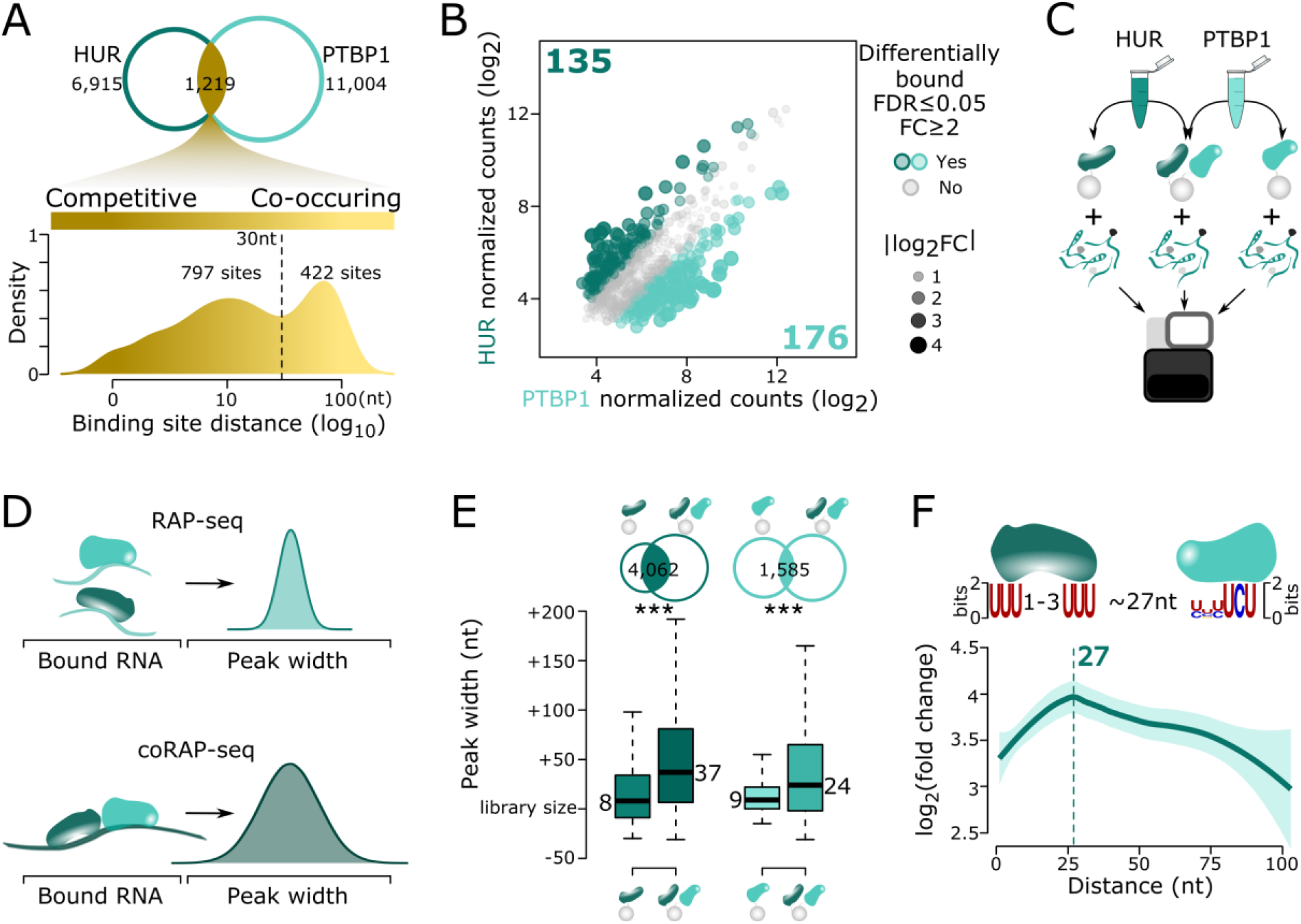
HUR and PTBP1 control RNA processing cooperatively with an optimal distance between their binding sites. **(A)** Venn diagram shows the overlap between HUR and PTBP1 peaks (top). Density plot (bottom) demonstrates the distance between peak summits of overlapping peaks. The x-axis represents the distance in nucleotides (nt) (log_10_ scale, bandwidth 0.15). The bimodal distribution is separated at 30 nt (dashed line). **(B)** Scatter plot compares HUR- and PTBP1-bound competitive peaks (dots, read count normalized, log_2_ transformed). Peaks either differentially bound by PTBP1 (light green) and HUR (dark green) or not differentially bound (grey) are highlighted (fold change ≥2, FDR-adjusted *p*≤0.05). Size and transparency of the dots are proportional to the absolute fold change. **(C)** Schematic representation of the co-RAPseq assay for HUR (dark green) and PTBP1 (light green) **(D)** Illustration of the expected effect on peak widths of cooperative binding events **(E)** Venn diagrams (top) show peak overlaps between the co-RAPseq with HUR (left) and PTBP1 (right) RAPseq assays, respectively. The intersections represent the binding sites used to assess the differences in peak widths. Boxplots (bottom) demonstrate differences in peak widths between the single- and co-RAPseq assays. The y-axis shows peak deviation from the median library size in nt per assay. Color codes are as in Figure 4C. Asterisks represent the significance of two-sided Wilcoxon rank sum tests between single- and co-RAPseq assays (***, p<0.001). **(F)** Model summarizing the cooperative binding of PTBP1 and HUR to their respective motifs (top). LOESS fitted regression line shows the trend in binding site enrichments (y-axis) when plotted as a function of the distance between the HUR and PTBP1 motifs (x-axis). The fold change in RNA binding is highest at 27 nt (dashed line).

To test cooperative binding properties of the two RBPs, we developed co-RAPseq, in which RNA binding is assayed in the presence of both proteins at the same time (**Figure 4C**). We hypothesized that in contrast to the single RBP assays, cooperative events determined by co-RAPseq lead to wider binding sites (peak width) due to the concomitant presence of the two proteins on the same RNA substrate (**Figure 4D**). While the peak width of the single RBP assays was 8 nt (HUR) and 9 nt (PTBP1) above median library size, co-RAPseq assays with both factors resulted in significantly wider 37 and 24 nt peak distributions, respectively (one-tailed Wilcox rank sum test, p<0.001) (**Figure 4E**). By measuring binding site enrichments in the co-RBP assay as a function of the distance between HUR and PTBP1 motifs, we determined that cooperativity was optimal at a 27 nt distance (**Figure 4F**). This optimal distance was neither resolvable in the single HUR and PTBP1 RAPseq assays nor when mapping co-RAPseq enrichment scores to the distances between motifs at control unbound sites (**Figure S4A-C**).

Of the 6,885 genes for which we detected binding events in either single or co-RAPseq assays, 873 were only present in the co-RAPseq assay (**Figure S4D**), suggesting that under steady-state conditions the post-transcriptional regulation of these gene products depended on the co-activity of both factors. In the single RAPseq assays, 86% (2,874/3,324) of the HUR and 59% (2,855/4,862) of the PTBP1-bound genes overlap with the co-RAPseq assay (**Figure S4D**). Of those genes, 19% (536/2,874 for HUR) and 22% (636/2,855 of PTBP1) were significantly stronger bound when both RBPs acted together (**Figure S4E**). In comparison to all 6,885 HUR- and PTBP1-bound genes, the uniquely and differentially bound genes in the co-RAPseq assay were functionally enriched in innate immune response pathways (**Figure S4F**) as previously observed for each factor individually (Christodoulou-Vafeiadou et al., 2018; Geng et al., 2021; Rothamel et al., 2021; Sasanuma et al., 2019; Sueyoshi et al., 2018). Remarkably, the strongest cooperative sites resided within introns of these pathways signifying cooperative functions of HUR and PTBP1 in mRNA pre-processing (**Figure S4G**).

### T7-RAPseq reveals m^6^A-dependent YTHDF1 binding to mediate mRNA regulation

To assess the transcriptome-wide relevance of RNA modifications in RBP-RNA recognition, we developed T7-RAPseq. We generated a modification-devoid RNA substrate by *in vitro* transcribing the native RNA substrate using T7 RNA polymerase (**Figure 5A**). When comparing RAPseq and T7-RAPseq, our results showed identical transcript distributions, high correlations (Spearman ρ=0.98 and Pearson r=0.97) (**Figure 5B**) and uniform read coverage over spliced and unspliced long and short transcripts (**Figure S5A**) confirming that the *in vitro* transcription step maintains substrate composition.

**Figure 5.**
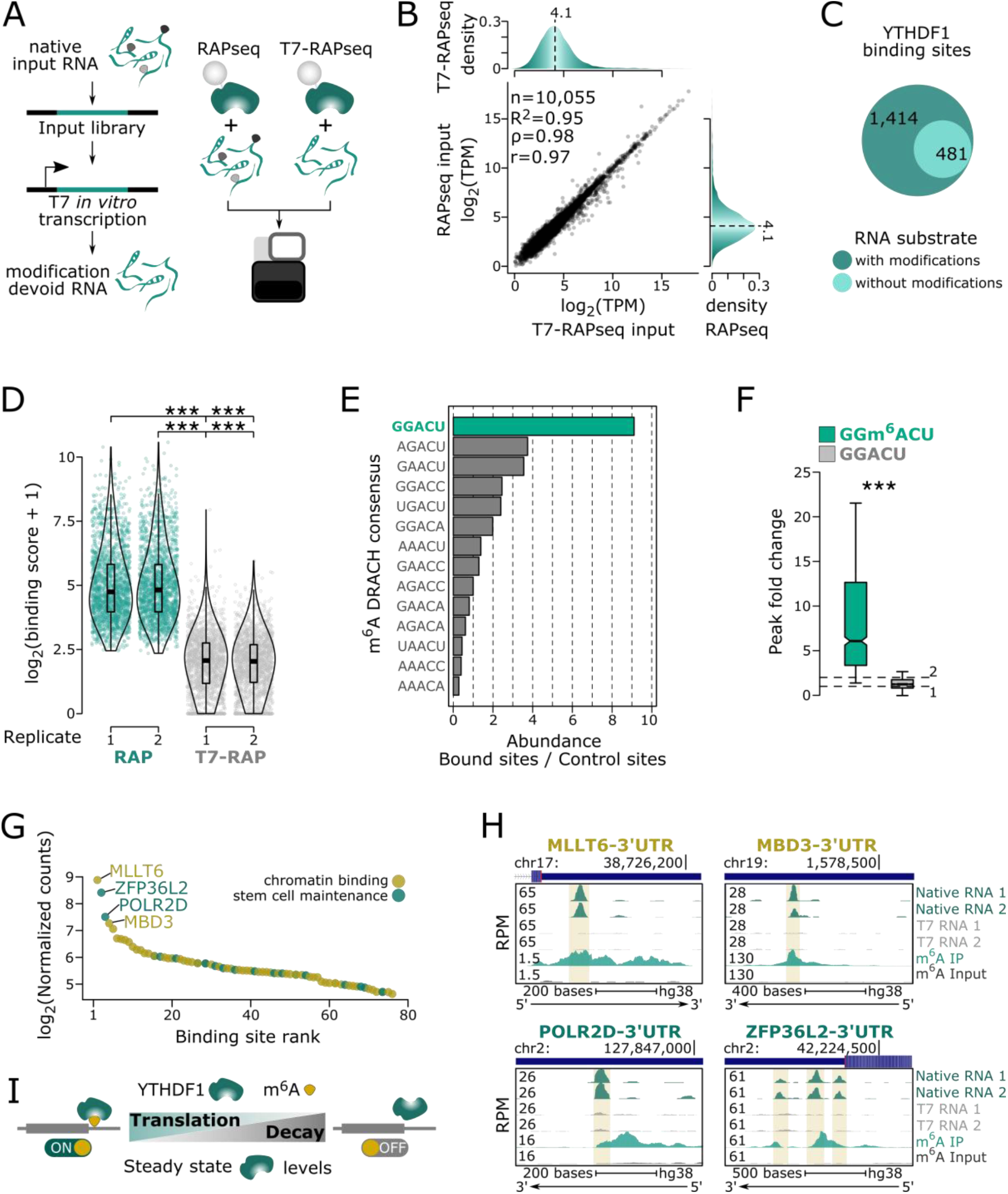
m^6^A is necessary for YTHDF1 to bind RNA at steady-state protein levels. **(A)** Schematic illustration of the T7-RAPseq method **(B)** Scatter plot compares the T7 *in vitro*-transcribed (T7-RAPseq) and the native RNA (RAPseq) substrates (log_2_ transformed transcripts per million, TPM). Numbers within the plot show Spearman (ρ), Pearson (r) correlation coefficient, linear regression (adjusted R^2^) and the number of observations. Density plots (right and top, bw=0.2) show the overall distributions of TPMs and their medians (4.1 log_2_ TPM, dashed line). **(C)** Venn diagram overlaps YTHDF1 binding sites between the native RAP and the T7-RAP substrates. **(D)** Violin and overlayed box plots show overall YTHDF1 enrichments (log_2_ transformed peak binding scores + 1) in both RAPseq (green points) with T7-RAPseq (grey points) replicates. Each point represents one binding site (n=1,895). Asterisks represent significance (Wilcoxon one-tailed rank sum test ***, p<0.001). **(E)** Barplot shows the frequency of DRACH 5-mers motif within YTHDF1 peaks normalized by their abundance at control sites. Top motif is highlighted (green). **(F)** Boxplots demonstrates the peak fold change of YTHDF1 enrichments in RNA substrates (n=1,059) with a modified (RAPseq, green) or modification depleted (T7-RAPseq, grey) GGACU motif. Dashed lines mark fold change values of 1 and 2. Asterisks represent the significance (Wilcoxon one-tailed rank sum test ***, p<0.001). **(G)** Dotplot illustrates ranking m^6^A-dependent YTHDF1 binding sites within mRNAs of genes belonging to two GO terms (Figure S5D). RAPseq binding site signal (library size normalized read counts log_2_ transformed, y-axis) is ranked (x-axis). Top 4 targets are highlighted. **(H)** Genome tracks show YTHDF1 binding to m^6^A within the 3’UTRs of the top 4 binding sites (MLLT6, MBD3, POLR2D and ZFP36L2) (Figure 5G). Genomic locations with scale bar indicate the length of the genomic region in bases and gene features (blue rectangle: exon and 3’UTR, arrow: direction of transcription) (x-axis) and normalized reads per million (RPM) (y-axis) are shown. Vertical yellow lines highlight bound site. Tracks for two replicates using native (dark green) or T7 *in vitro* transcribed (grey) input RNA as well as m^6^A-specific RNA immunoprecipitation (IP, light green) and the respective input control are presented. **(I)** Model illustrates m^6^A-dependent roles of YTHDF in post-transcriptional regulation. The presence of m^6^A (yellow circle) enables YTHDF1 (green reniform cartoon) binding to its RNA targets (grey box) and thereby enhances their stability and translation under steady state conditions. In contrast, the absence of m^6^A drastically abolishes RNA binding of YTHDF1 resulting in RNA decay and decreased translation.

We applied T7-RAPseq to determine modification-dependent targets of the m^6^A binder YTHDF1 (X. Wang et al., 2015). According to previous reports (X. Wang et al., 2015), YTHDF1 binds to a preferred GRAC (R is an A or G) motif of the m^6^A DRACH consensus sequence (D is A, U or G and H is A, U or C). About 75% (1,414/1,895) of the identified YTHDF1 binding sites were strictly dependent on the presence of m^6^A (**Figure 5C, Table S4**) and the binding affinity drastically decreased in the absence of RNA modifications (one-tailed Wilcox rank sum test, p<0.001) (**Figure 5D**) corroborating m^6^A dependency in RNA binding. We observed that GGACU was the most abundant m^6^A-DRACH motif (**Figure 5E**) either as a 5- or its two 4-mers GGAC and GACU (**Figure S5B**). Although enrichment scores for the pentamer were higher than for the individual tetramers (one-tailed Wilcox rank sum test, p<0.001) (**Figure S5C**), the enrichment scores are comparable when the two tetramers can act cooperatively (one-tailed Wilcox rank sum test, p<0.001) (**Figure S5C**). This demonstrated that GGACU is the optimal binding motif and the two suboptimal tetramers (GGAC and/or GACU) can increase the frequency of YTHDF1 interactions with RNA and/or induce the formation of YTHDF1 dimers. In accordance with the m^6^A dependency of YTHDF1 (Xu et al., 2015), GGACU pentamer containing binding sites with a m^6^A had higher enrichment scores when compared to m^6^A depleted motifs (one-tailed Wilcox rank sum test, p<0.001) (**Figure 5F**).

We next investigated pathways that could be affected by the presence or absence of YTHDF1 m^6^A binding. We found 48 significantly enriched (FDR-adjusted *p*<0.01) parent GO terms with the highest m^6^A dependency in genes involved in stem cell maintenance and chromatin binding (**Figure 5G** and **S5D**). We used public m^6^A-sequencing data to validate the top 4 binding sites in MLLT6, ZFP36L2, POLR2D and MBD3 mRNAs (**Figure 5G-H**) (Dominissini et al., 2012). In conclusion, the deterministic difference in RAPseq and T7-RAPseq data allowed to discern a clear m^6^A-dependent on-off switch for YTHDF1 binding (**Figure 5I**) and that T7-RAPseq can be used to distinguish genome-wide contributions of RNA modifications to RBP binding activities.

### Non-canonical RBPs have specialized RNA interactomes

Despite the increased identification of ncRBPs through latest technological advances, knowledge about the modes of ncRBP-RNA binding is limited. In order to test whether RAPseq can discover RNA targets of ncRBPs, we intersected RBPomes identified in four independent studies (Castello et al., 2012; Queiroz et al., 2019; Trendel et al., 2019; Urdaneta et al., 2019). Across all assays, one third (243/718) of all common RBPs were ncRBPs (**Figure 6A**). We selected 26 ncRBPs containing different domain composition and canonical functions (**Table S1**). While cRBPs bound on average to 9,574 sites, we observed two groups of ncRBPs binding to either more or fewer then 1000 sites using RAPseq. Ten ncRBPs had a mean of 1,757 (range: 1,102-3,134) and the remaining 16 on average 113 (range: 7-534) binding sites (**Figure S6A**). PEBP1 had the highest (3,134) and CCDC124 (7) the lowest number of binding sites (**Figure S6B**). While cRBPs had a median of 2.5 (range: 2.1-3.2) binding sites, ncRBPs had one binding site across the gene body (**Figure S6C**).

**Figure 6.**
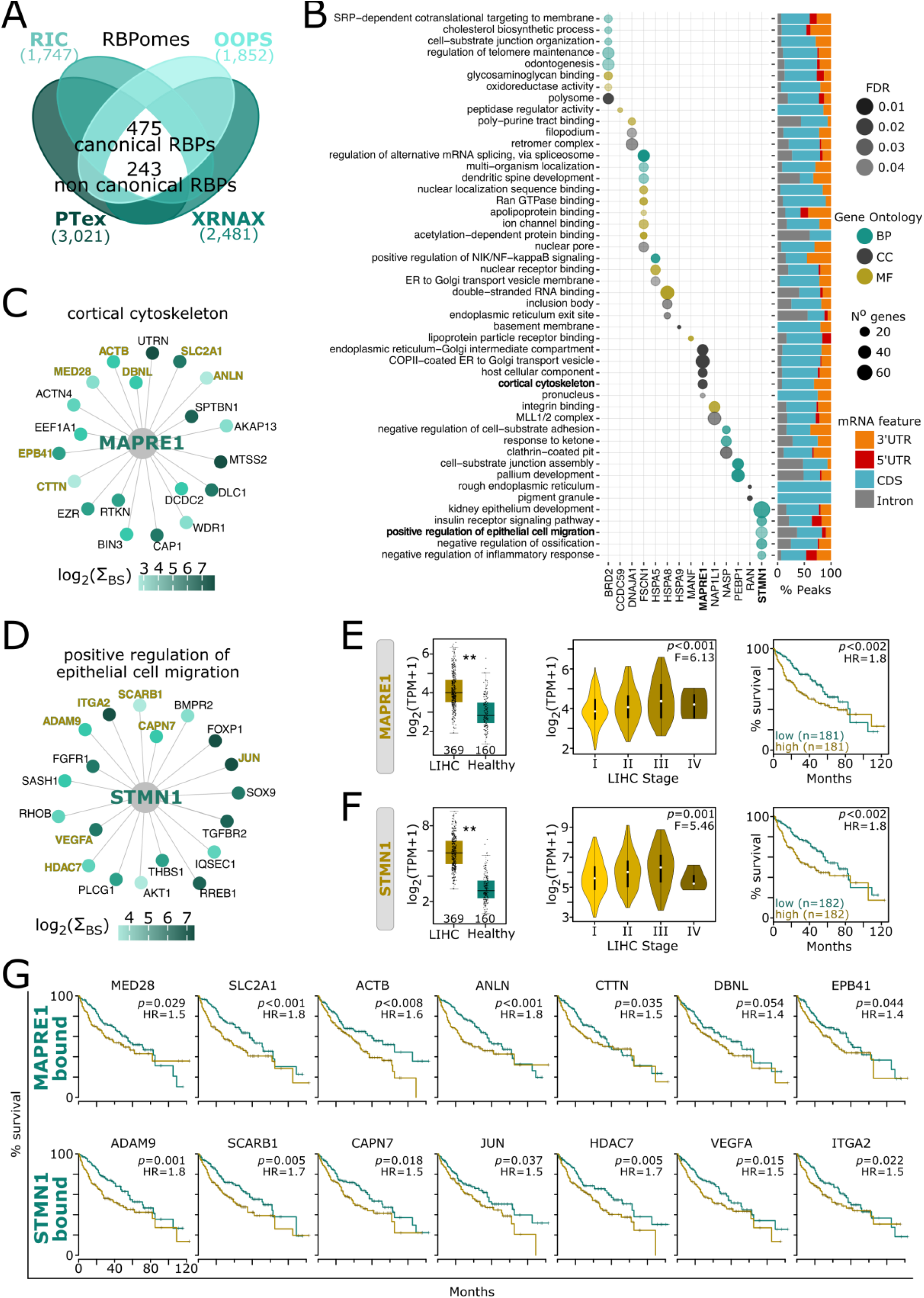
ncRBPs bind to cancer-associated transcriptomes. **(A)** Four-way Venn Diagram overlaps reported RBPs identified by RNA interactome capture (RIC), orthogonal organic phase separation (OOPS), protein and phenol-toluol extraction (PTex) and cross-linked RNA extraction (XRNAX). The number of RBPs discovered by each method is written in parenthesis. Common RBPs (white) are divided into canonical or noncanonical. **(B)** Circle plots (left) display ncRBP-RNA interactomes functionally enriched in gene ontology (GO) terms. Circle diameter is proportional to the number of enriched genes per category; circle color corresponds to the type of GO term (BP: biological process, CC: cellular compartment, MF: molecular function) and circle transparency to FDR-adjusted *p*-values. Stacked barplot (right) shows the proportional binding to mRNA features (color-coded) for each GO term. (**C, D**) Interaction network shows functionally enriched (**C**) MAPRE1- and (**D**) STMN1-RNA interactomes. Nodes represent gene names of MAPRE1 and STMN1-bound mRNAs and color displays peak binding scores intensities for all binding sites (sum log_2_ transformed; light blue: low, dark blue: high) mapping to the gene representing the node. Highlighted genes (yellow) are further studied in **G**. (**E, F**) Liver Hepatocellular Carcinoma (LIHC) association of (**E**) MAPRE1 and (**F**) STMN1. Boxplots (left) show MAPRE1 and STMN1 expression levels (log_2_ TPM+1) in LIHC patients (red) and healthy human liver tissues (green) from TCGA and GTex datasets. Asterisks indicate *p*-values (one-way ANOVA F test, **p<0.01). Violin plots (middle) show increasing MAPRE1 and STMN1 expression levels with LIHC stage severity. Linear regression and *p*-values (one-way ANOVA F test) are noted. Kaplan-Meier plots (right) show association of MAPRE1 and STMN1 low (green) and high (yellow) expression levels and ten-year survival within the LIHC cohort. Low and high groups are separated by median gene expression level. Patient number (n) per group, hazard ratios (HR) and *p*-values (Mantel-Cox test) are reported. (**G**) Kaplan-Meier plots display an association between expression levels of genes highlighted in panels **C** and **D** (yellow) and overall survival (as in **E** and **F**, right). Corresponding hazard ratios (HR) and *p*-values (Mantel-Cox test) are shown (n=180-183 patients for each high and low group).

By determining interactions between ncRBPs and RNA classes, we found that most binding sites (median: 97.2%, range: 60-100) resided in mRNAs. Compared to the other ncRBPs, RAN exhibited preferences for exonic regions and PRDX6 for 5’UTRs. CCT2 and TKT had the highest proportional enrichment for long and HSPA5 for small noncoding RNAs (ncRNAs) (**Figure S6D**). In comparison to cRBPs, ncRBPs exhibited weaker binding strengths (**Figure S6E**). HSPA8 and PEBP1 displayed both the highest number of binding sites and the strongest enrichments (**Figure S6C and S6F**). These results implied more specialized functions as post-transcriptional regulators as well as a higher dependency on regulatory signals and interactions due to less *bona fide* functions in competitive RNA binding.

To assess the specialization for each ncRBP-RNA interactome, we performed a functional gene enrichment analysis of the bound mRNAs. For 14 ncRBPs, we identified 48 GO terms that were significantly and specifically enriched (FDR-adjusted *p*≤0.05) (**Figure 6B**). GO terms with the highest percentage of binding sites in 5’ and 3’UTRs were obtained for genes enabling apolipoprotein binding (for FSCN1) and negative regulation of inflammatory response (for STMN1). In addition, we found mRNA enrichments for genes in cortical cytoskeleton (for MAPRE1) and epithelial cell migration (for STMN1). The canonical function of these two ncRBPs is to either promote (MAPRE1) or destabilize (STMN1) microtubule elongation (**Figure 6B-D**) (Askham et al., 2002; Steinmetz et al., 2000; van Haren et al., 2018; Wittmann et al., 2004). Since we profiled the MAPRE1- and STMN1-RNA interactomes in liver cancer cells (HepG2), we inspected the association with hepatocellular carcinoma (LIHC). MAPRE1 and STMN1 have been linked to promote liver cancer progression through mechanisms exerted by the canonical functions in cytoskeletal remodeling (Aiyama et al., 2020; Lu et al., 2018; Orimo et al., 2008; Wong et al., 2008). We found that upregulation of MAPRE1 and STMN1 gene expression correlated with the severity of the pathological stage and significantly decreased patient survival in LIHC patients (**Figure 6E-F**). A subset of MAPRE1 and STMN1 RNA targets (**Figure 6C-D**) was significantly associated with poorer patient survival in LIHC (**Figure 6G**). Thus, as post-transcriptional regulators, MAPRE1 and STMN1 could increase LIHC progression by either enhancing stability and/or localized translation of their mRNA targets.

### Cancer-associated variants of the IGF2BP family show altered RNA binding

The family of insulin-like growth factor-2 mRNA binding proteins (IGF2BP) comprises three paralogs (IGF2BP1, IGF2BP2 and IGF2BP3). In human and mouse, IGF2BP genes show oncofetal expression patterns with high expression during embryonic development and in liver cancer (**Figures S7A-C**) (Cardoso-Moreira et al., 2019; Rudolph et al., 2016). We inspected frequently occurring IGF2BP variants in cancer types and found one common amino acid change in IGF2BP1 (R167C/H) and two in IGF2BP3 (I474M and R525C) (**Figure 7A**). Frequent missense mutations were absent in IGF2BP2 but a cancer-associated splicing variant (isoform B) (**Figure 7A**) was previously identified (Tybl et al., 2011; J.-Y. Zhang et al., 1999). We generated IGF2BP paralogs (wt) and cancer-associated variants to compare their RNA binding capacities by RAPseq (**Figure 7B**). The paralogs showed highly similar binding preferences towards CA-rich motifs, as previously reported (Conway et al., 2016; Schneider et al., 2019). We found that these motif preferences were maintained in the cancer-associated variants (Spearman rank correlations, median=0.96, range: 0.92-0.99, n=28) (**Figure S7D**). In contrast, k-mer enrichment profiles differed when comparing IGF2BP paralogs and variants to either RBFOX2 or YTHDF1 (**Figure S7D**). Compared to the IGF2BP1 paralog, the cancer-associated variant IGF2BP1-R167C showed similar but IGF2BP1-R167H had overall lower signals (**Figure 7C**). We found no global differences between the two IGF2BP2 isoforms. IGF2BP3 variants exhibited weaker binding strengths when compared to the wt (**Figure 7C**). By testing differential RNA binding, we found no differences between the wt and the IGF2BP1-R167C variant but the wt bound to significantly more sites (n=962) than the IGF2BP1-R167H variant (n=34). The two IGF2BP2 variants had a small and balanced number of differently bound sites (56 for isoform B and 70 for isoform A). IGF2BP3 bound to more sites than the two mutants (268 *versus* 15 for IGF2BP3-I474M and 102 *versus* 3 for IGF2BP3-R525C) (**Figure 7D**). We noticed that the differentially bound sites of IGF2BP2 isoform B occurred more frequent in ncRNAs than those of isoform A (Chi-squared test, p<0.003) (**Figure 7E**). Thirteen ncRNAs were more bound by isoform B compared to only four by isoform A and the highest differences were observed for snoRNAs and miRNAs (**Figures S7E-F**).

**Figure 7.**
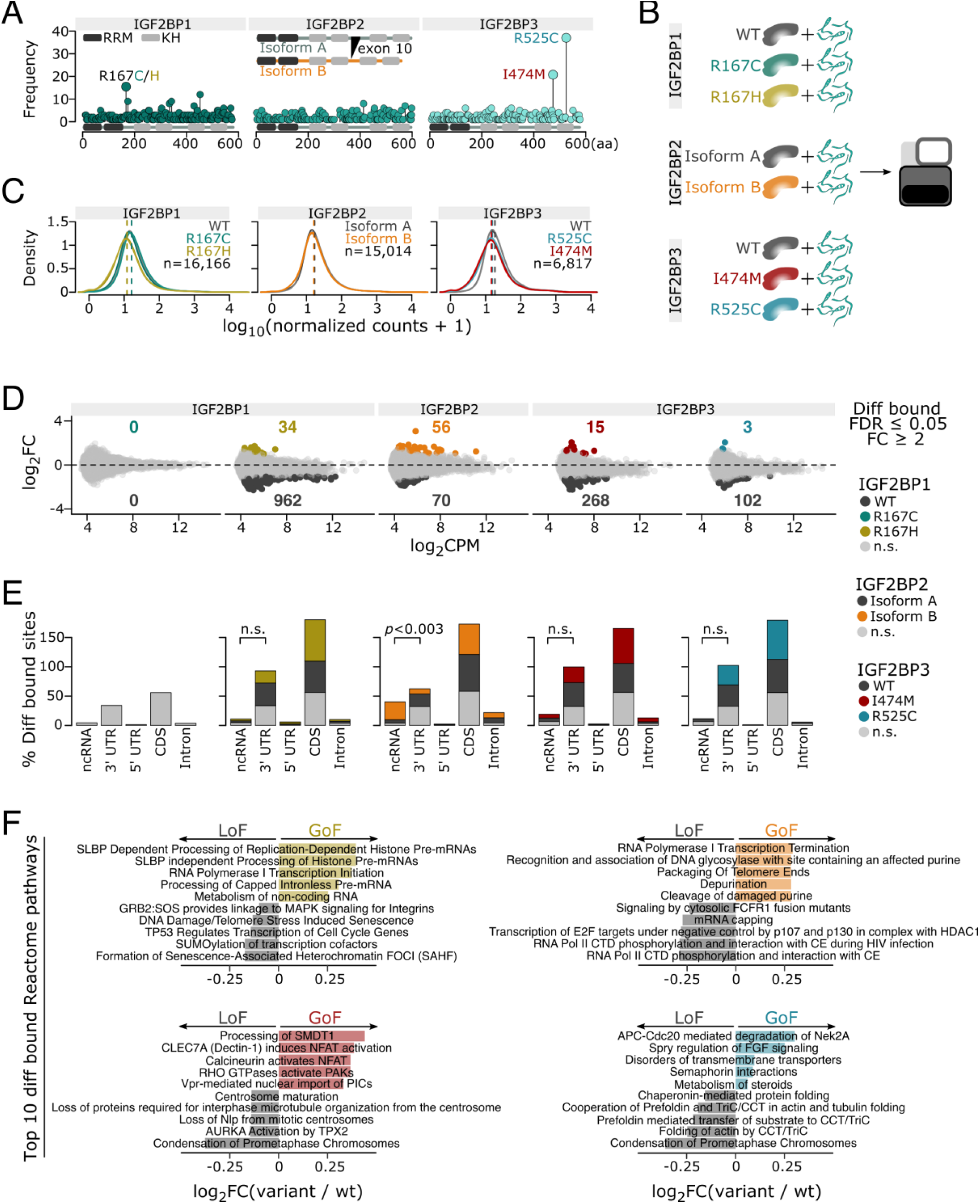
Pathological variants of the IGF2BP family cause functional gains and losses in RNA binding. **(A)** Bars illustrate RRM and KH domain locations in each paralog protein (length in amino acids, aa, x-axis). Lollipops show aa mutation frequencies (y-axis) in cancer patients. Mutants selected for RAPseq profiling are color-coded. The pathogenic isoform B of IGF2BP2 with the skipped exon 10 is drawn. **(B)** Schematic representation of RAPseq assays performed to profile IGF2BP paralogs and pathological variants (color-coded). **(C)** Density plots represent the distribution of RAPseq signals (read counts normalized to HaloTag in log_10_) for each IGF2BP variant with respect to their wild types. Dashed lines indicate medians. The number of observations (n) is shown in each panel. **(D)** MA plots demonstrate differentially bound sites (DBS) between wild type and variants (y-axis) as a function of the average signal per binding site (x-axis, in log_2_ counts per million). Data points and number represent significantly (color-coded by variant) and not significantly DBS (grey) sites (fold change ≥2, FDR-adjusted *p*≤0.05). **(E)** Stacked barplots show the percentage of DBS (y-axis) binned per major RNA feature for each wild type and variants (color-coded) (x-axis). Bars of the same color sum up to 100%. Significant difference of DBS in 3’UTR and ncRNA of the two IGF2BP2 isoforms (Chi-squared test with continuity correction *p*-value) are highlighted. **(F)** Horizonal barplots display the top 10 differentially bound Reactome pathways between variants and wild types (color-coded, fold change ≥2, FDR-adjusted *p*≤0.05).

At the pathway level, we observed significant differences (FDR-adjusted *p*≤0.05) between variants and respective wt paralogs despite the small size effects (**Figure 7F**). The IGF2BP1-R167H variant showed a slight increase in binding towards pathways regulating RNA metabolism and a decrease in binding to factors involved in DNA damage response. This could suggest a gain of function (GoF) shift towards promotion of growth and loss of function (LoF) in regulation of factors safeguarding genotoxic events. For the pathological IGF2BP2 isoform B, we observed a GoF in DNA damage response and a LoF in growth promoting pathways. For IGF2BP3-I474M, we noticed LoFs in chromosome and centrosome organization during mitosis, which as for IGF2BP1-R167H could result in genotoxic events plausibly through missegregation during anaphase and a GoFs in calcium-dependent overproduction of ATP and growth promotion. In the IGF2BP3-R525C mutant, a GoF controlled binding to growth and to mobility promoting mRNAs as well as a LoF affecting binding of the chaperonin complex CCT/TriC. Overall, our data suggested that the profiled mutations alter binding strength but not binding specificities, which can result in post-transcriptional deregulation of their RNA targets and related pathways.

## DISCUSSION

We have developed RAPseq, a method that provides *in vitro* derived RBP-RNA interactomes by using recombinant RBPs and native cellular RNA substrates with endogenous epitranscriptomic modifications maintained. RAPseq can be applied in versatile and large-scale transcriptome-wide studies to address fundamental questions in post-transcriptional gene regulation.

First, RAPseq allowed us to design combinatorial cross-species RBP-RNA interaction assays to dissect vertebrate conservation of HUR’s binding specificities and the divergence of its intronic and 3’UTR binding sites between lower and higher vertebrates. Second, we used RAPseq to identify parameters for optimal cooperativity between HUR and PTBP1 and pathways that benefit from this co-binding activity. Third, we produced a modification depleted version of the native RAPseq substrate (T7-RAPseq) and discovered which YTHDF1 binding sites depend on the presence of m^6^A. Forth, we demonstrated that RAPseq can overcome one of biggest limiting factors in current RNA binding assays by multiplexing the profiling of 26 ncRBP-RNA interactomes. We concluded that ncRBPs have specialized rather than global post-transcriptional regulatory roles and found that some moonlighting activities coincided within the same biological processes described for the canonical functions. Fifth, we profiled cancer-associated variants of the IGF2BP family. Except for IGF2BP2, the other variants bound to a larger extent to RNA substrates involved in pathways promoting cell growth. This suggested that increased RNA binding activities of frequently mutated IGF2BPs could provide proliferative advantages in cells of cancer patients.

We showed that RAPseq can be used by individual laboratories to enable consortia-like studies by describing applications of the highest interest. Additional method developments, including co- and T7-RAPseq, extended the spectrum of practical implications. Increasing the proportion of currently underrepresented intronic sequences (**Figure S2G**) in the RAPseq substrate, i.e., by purifying RNA from nuclear isolates will be of relevance in the future to assess alternative splicing events that have co-RBP regulatory dependencies. The increasing number of ncRBPs without known RNA binding functions but with well-established roles in cellular metabolism, opens questions regarding contextuality of their RNA binding activity. For example, binding to RNA and executing known functions could occur simultaneously or may be mutually exclusive. RNA-enzyme-metabolite (REM) networks (Hentze & Preiss, 2010) proven so far, have demonstrated a few examples of linking interdependent functions. RAPseq could be used to perform RNA binding assays in the presence or absence of such metabolites across cell type-specific transcriptomes varying in composition to assess increased or disrupted interdependency of REM networks.

Altogether, RAPseq is a versatile and above all enabling tool that will ease and pave the way towards the description of the complete RBPome and help envision new types of studies that would be unfeasible otherwise.

## Acknowledgments

This work was supported by KI KID KI-KID funding (2016–00189, CK), Knut & Alice Wallenberg foundation (KAW 2016.0174, CK); Ruth & Richard Julin foundation (2017– 00358, 2018–00328, 2020-00294 CK); SFO-SciLifeLab fellowship (SFO_004, CK); Swedish Research Council (2019-05165, CK); Lillian Sagen & Curt Ericsson research foundation (2021-00427, CK); Gösta Milton’s research foundation (2021-00527, CK); Onassis Foundation (F ZO 057-1/2018-2019, SP), Karolinska Institute (2019-01080 and 2019-01102, IA), Erasmus+ (20170801, IA; 20200115, NP); Swedish National Infrastructure for Computing (SNIC) at UPPMAX (SNIC 2017/7-260, uppstore2018110, SNIC 2020/15-293, SNIC 2020/16-228, CK-IA). We would like to thank the National Genomics Infrastructure in Stockholm funded by Science for Life Laboratory, the members of the Marc Friedländer, Vicent Pelechano, and Claudia Kutter laboratories as well as Carsten Daub for helpful feedback.

## Author contribution

CK conceptualized the study. IA and CK designed experiments and computational analysis. IA, SP and NP performed experiments. IA and SP performed computational analysis. CK acquired funding. IA and CK wrote the manuscript. All authors read and approved the final manuscript.

## Supplementary Figures

**Figure S1.**
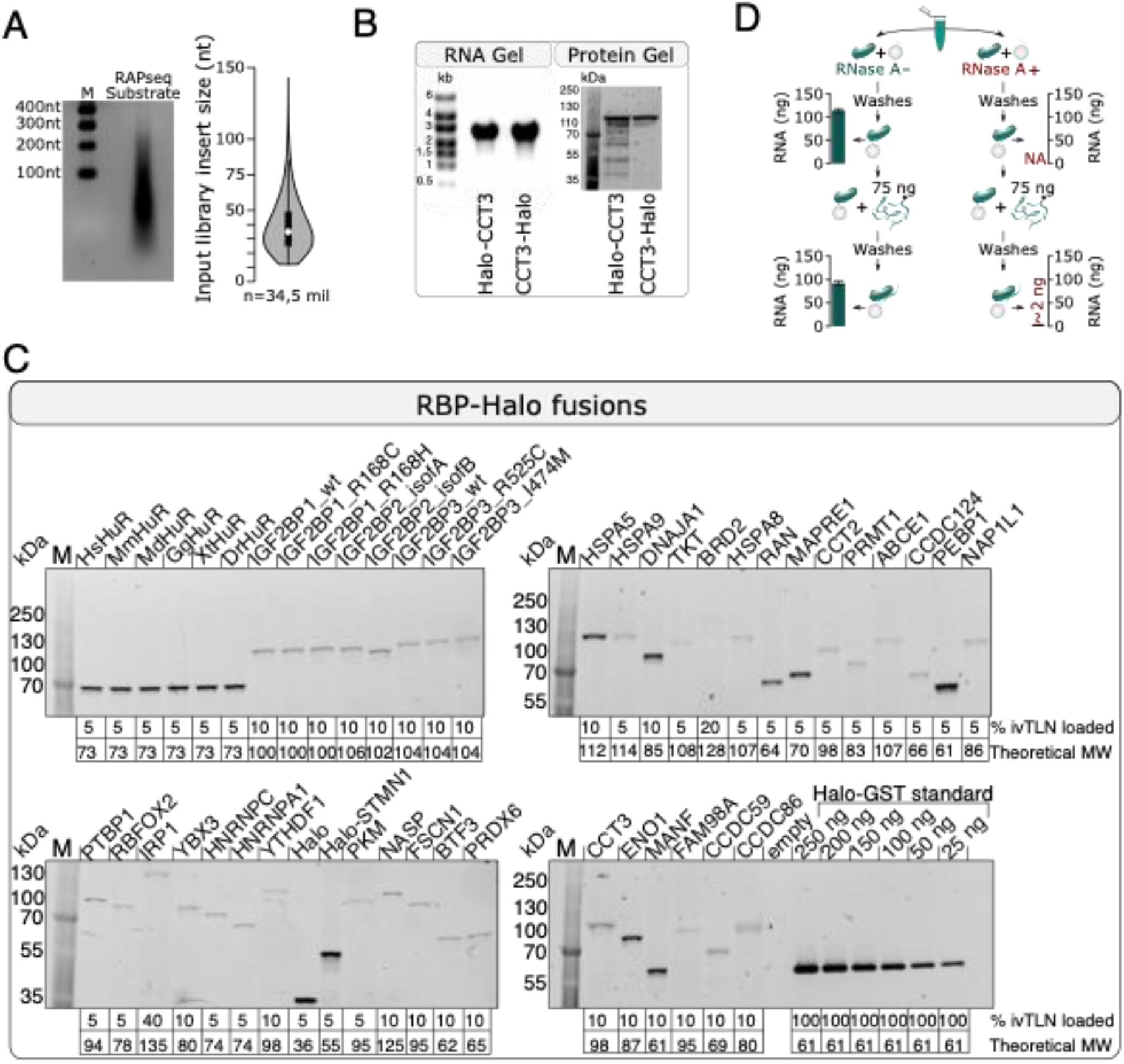
RBPs are produced as C-terminal Halo fusions and the RAPseq substrate is produced by random fragmentation. Related to Figure 1. **(A)** Agarose gel (left) and violin plot (right) show size distribution of the sequenced input library (34.5 million reads). **(B)** Agarose (left) and SDS polyacrylamide gel (right) represent *in vitro* transcribed RNA (left) and *in vitro* translated protein products of N- and C-terminal RBP-HaloTag^®^ fusion products resulting in truncated or full-length protein products, respectively. Protein products are visualized by HaloTag^®^ bound to Halo Alexa488-Ligand before gel electrophoresis. N-terminal tagged CCT3 displays a mix of full length and truncated translation products. **(C)** SDS-PAGE gels of all described RBP-Halo fusion proteins. The percentage of loaded protein refers to the amount of *in vitro* translation (ivTLN) reaction used for the RAP binding assay. Theoretical RBP molecular weight (MW) in kDa is reported. A recombinant Halo-GST fusion protein was used as a titration curve. **(D)** Illustration and bar plots confirm the amount of RNA recovered before (top) or after (bottom) the RAP binding assay using *in vitro* translated HUR-Halo fusion proteins in either the absence (left) or presence (right) of RNase A (barplots, error bars show ± SEM of three replicates). The eluted RBP bound RNA generally ranges from 1 to 5 ng depending on the RBP, higher amounts are indicative of contaminants from the *in vitro* transcription and translation.

**Figure S2.**
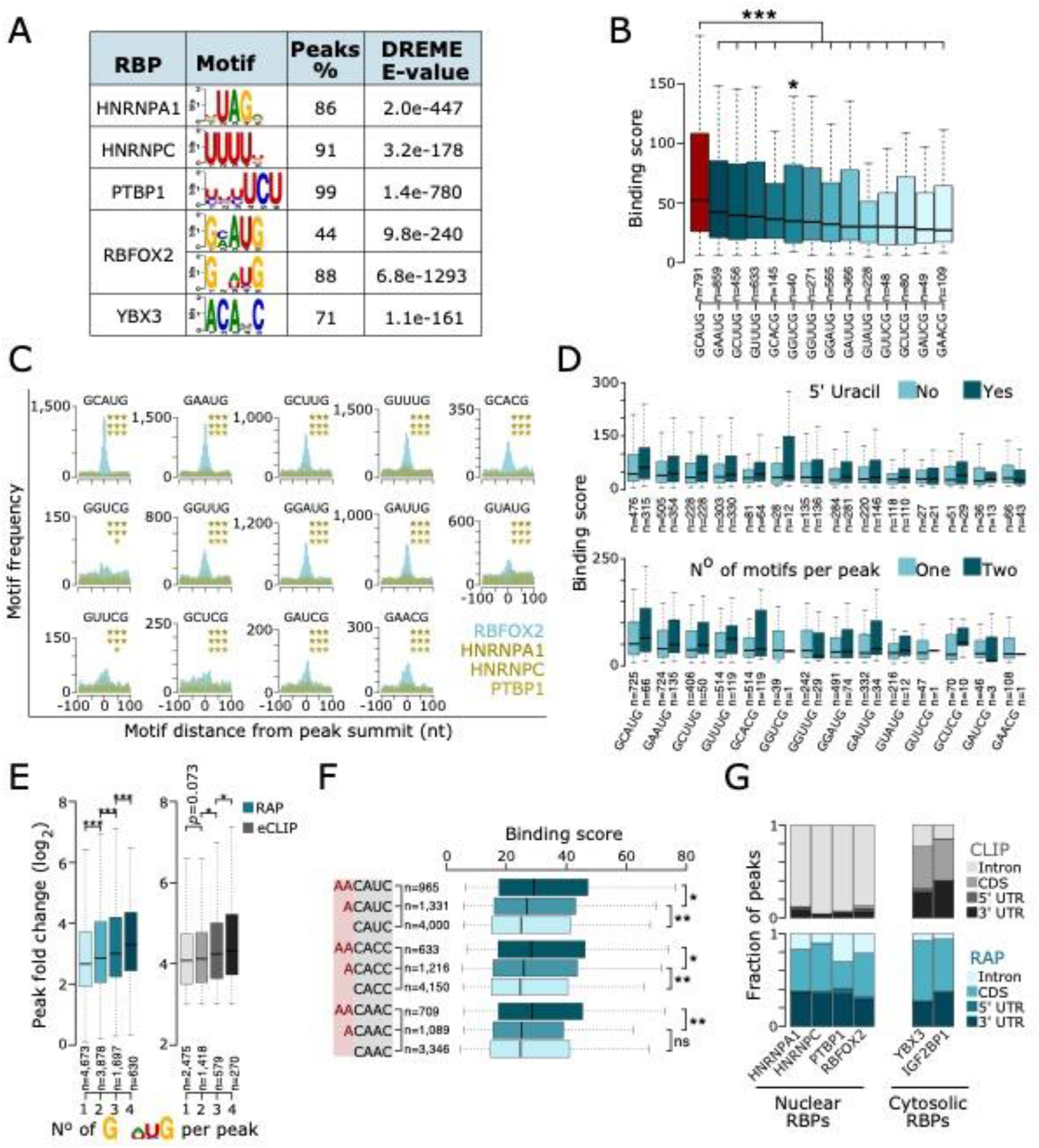
RAPseq confirms and identifies new secondary motifs. Related to Figure 2. **(A)** Table lists the percentage and corresponding DREME algorithm E-values of motifs in the RBP-bound transcriptome. **(B)** Boxplots show peak binding scores and number (n) of peaks containing one of the 14 RBFOX2 ‘GNWYG’ motifs. Asterisks represent *p*-values (one-tailed Wilcoxon rank sum test, *p<0.05, ***p<0.001). **(C)** Metaplots compare the frequency of the 14 RBFOX2 ‘GNWYG’ motifs (polygons) with respect to the RBFOX2 (light blue) and three control RBPs (HNRNPA1, HNRNPC and PTBP1) (yellows) peak summits (± 100 nt, bin size 5 nt). Asterisks represent *p*-values (binomial test for differential central enrichment in RBFOX2 to the three control RBPs, *p<0.05, ***p<0.001. **(D)** Boxplots display peak binding scores as a function of the presence of an additional 5’ uracil at each GNWYG 5-mer (top) and as a function of number of 5-mers per peak (bottom). **(E)** Boxplots show peak fold changes as a function of number of RBFOX2 ‘GNWYG’ motif occurrences in RAPseq (blue) and eCLIPseq (grey) assays. Asterisks represent *p*-values (one-tailed Wilcoxon rank sum test, *p<0.05, ***p<0.001). **(F)** Boxplots display peak binding scores as a function of the presence of one or two additional 5’ adenosines at the three 4-mers of the YBX3 ‘CAHC’ core motif. **(G)** Stacked barplots represent the fraction of peaks for six RBPs residing in a mRNA feature in RAPseq (blue) and eCLIPseq (grey) assays.

**Figure S3.**
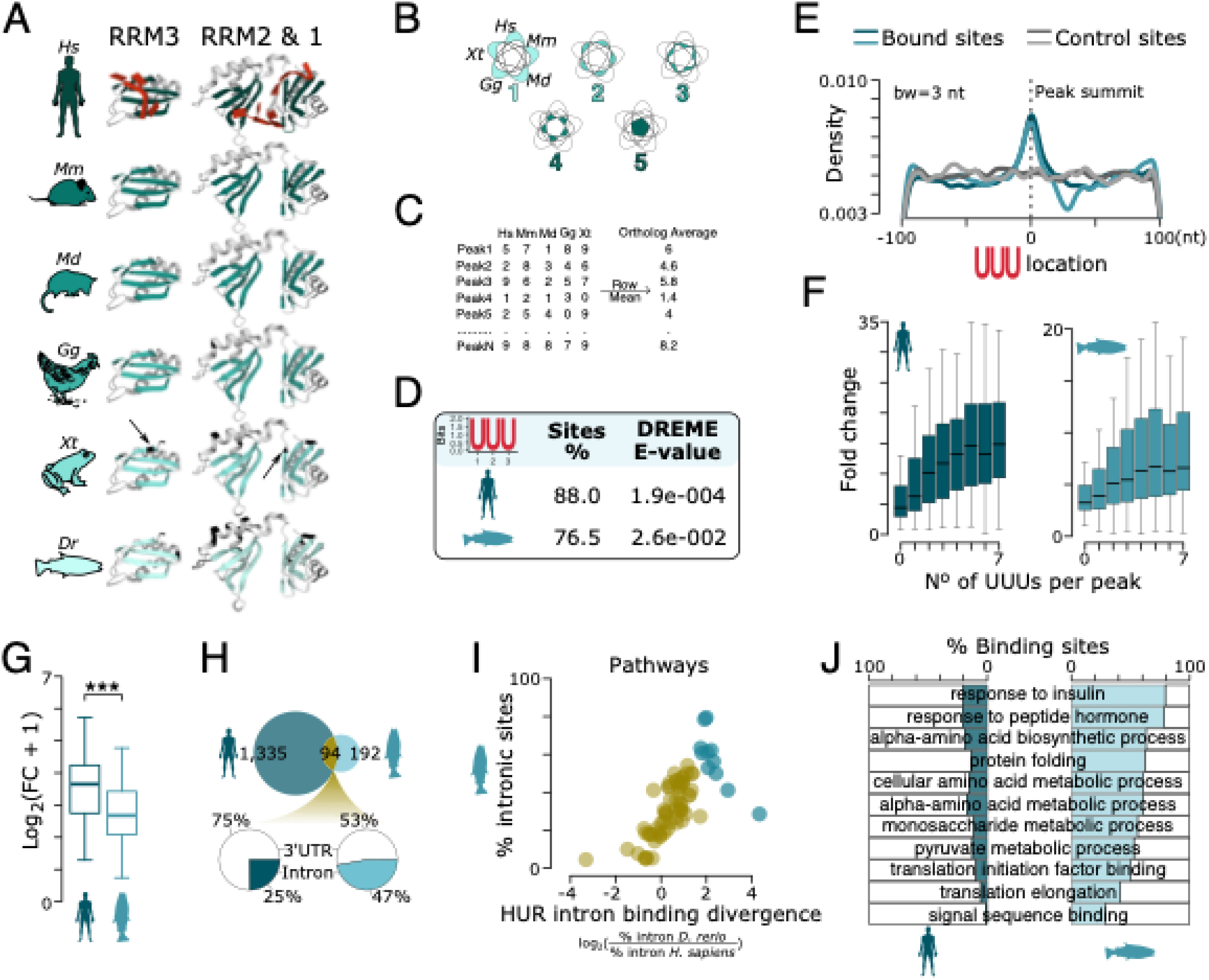
Combinatorial cross-species RAPseq assays dissects HUR conserved and divergent roles. Related to Figure 3. **(A)** Crystal structures highlight amino acid differences (black arrows) of HUR orthologs in beta sheets (green) or alpha helices (grey) in the RRM1 to 3. The human HUR is in complex with RNA (uracils in red). **(B)** Schematic representation of intersections used in Fig. 3G-I (x-axes). **(C)** Table exemplifies calculations of ortholog average scores. **(D)** Table lists the percentage of peaks with an uracil triplet (±50 nt around the peak summit) and corresponding DREME E-value of for the human and fish HUR orthologs in RAPseq assays with the fish substrate. **(E)** Line plot (smoothing band width 3 nt) shows frequency of tetrapod HUR binding to uracil triplets ± 100 nt of the peak summit (dark blue, human; light blue, zebrafish) and control sites residing 1000 nt distant from the peak summit (dark grey, human; light grey, zebrafish). **(F)** Boxplots display peak fold changes by number of uracil triplets ± 100 nt of the peak summit (human: dark blue, zebrafish: light blue). **(G)** Boxplot shows peak fold changes over all binding sites of human (n=3,191) and zebrafish (n=2,975) HUR using zebrafish RNA. Asterisks represent *p*-values (one-tailed Wilcoxon rank sum test, ***p<0.001). **(H)** Venn diagram (top) overlaps GO terms of genes bound by human and zebrafish HUR using zebrafish RNA. Pie charts (bottom) display the percent of binding sites within introns and 3’UTRs of genes in the overlapping GO terms (yellow) in human (left) and zebrafish (right). **(I)** Scatter plot shows the percent of intronic HUR binding sites with respect to 3’UTR for genes in each GO term as a function of the ratio between the percentage of zebrafish and human intronic sites. Each point represents one overlapping GO term (94, yellow) with the top intronic binding site bearing GO terms highlighted (blue). **(J)** Bar plots (left, human; right, zebrafish) show the percent of binding sites within introns (dark blue, human; light blue, zebrafish) and 3’UTRs (white) of GO terms highlighted (blue) in Figure S3I. GO terms are ranked based on the percentage of intronic sites in zebrafish.

**Figure S4.**
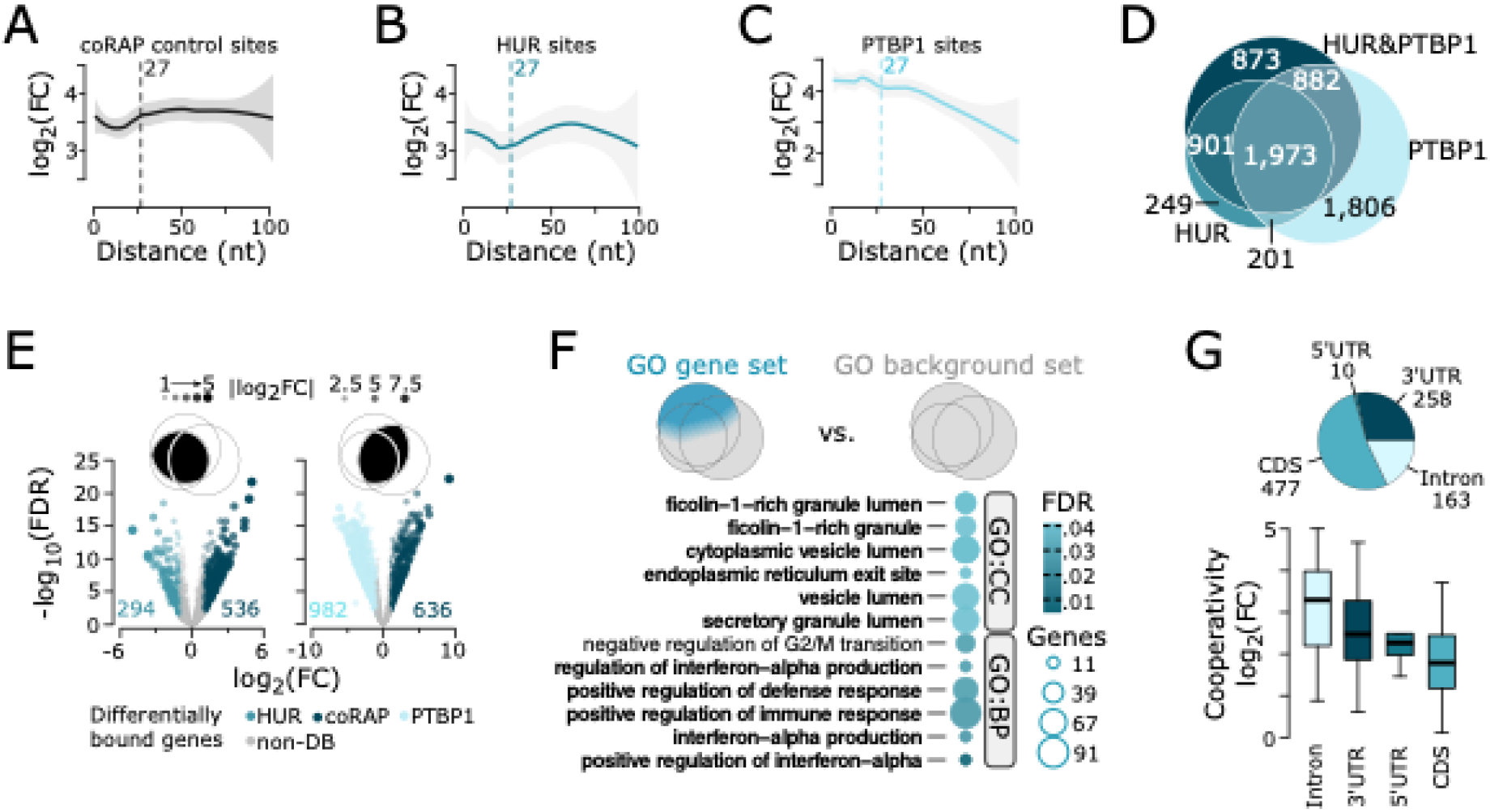
HUR and PTBP1 cooperate to regulate interferon response pathways and secretory processes in HepG2 transcriptomes. Related to Figure 4. **(A-C)** LOESS fitted regression lines show the trend in binding site enrichments (y-axis) when plotted as a function of the distance between the HUR and PTBP1 motifs (x-axis). Enrichment scores are mapped to the distance of adjacent unbound control sites in (**A**) HUR and PTBP1 co-RAPseq, (**B**) single HUR and (**C**) PTBP1 RAPseq assays. **(D)** 4-way VENN diagram overlaps genes between co- and single RAPseq assays. **(E)** Volcano plots demonstrate differentially bound (DB) genes between co-RAPseq *versus* HUR (left) and PTBP1 (right) RAPseq. VENN diagrams (top) illustrate which genes were tested. Genes either differentially bound in the co-RAPseq (dark blue), HUR (blue), PTBP1 (light blue) RAPseq assay or non-DB (grey) are highlighted (fold change ≥2, FDR-adjusted p≤0.05). Number of DB genes are mentioned. Size and transparency of the dots are proportional to the absolute fold change. **(F)** Circle plot displays gene ontology (GO) terms enriched for genes bound only and more in the co-RAPseq assay highlighted in blue in the 3-way VENN diagram (top) using all other genes identified as background gene set. Circle diameter is proportional to the number of enriched genes per GO term (CC: cellular compartment, BP: biological process) and color intensity to FDR-adjusted *p*-values. **(G)** Pie chart (top) illustrates the distribution of the binding sites to mRNA features in the GO terms listed in Figure S4F. Boxplots (bottom) show fold-change of binding site enrichments for each mRNA feature. Number of binding sites mapping to each mRNA feature (color-coded) are indicated.

**Figure S5.**
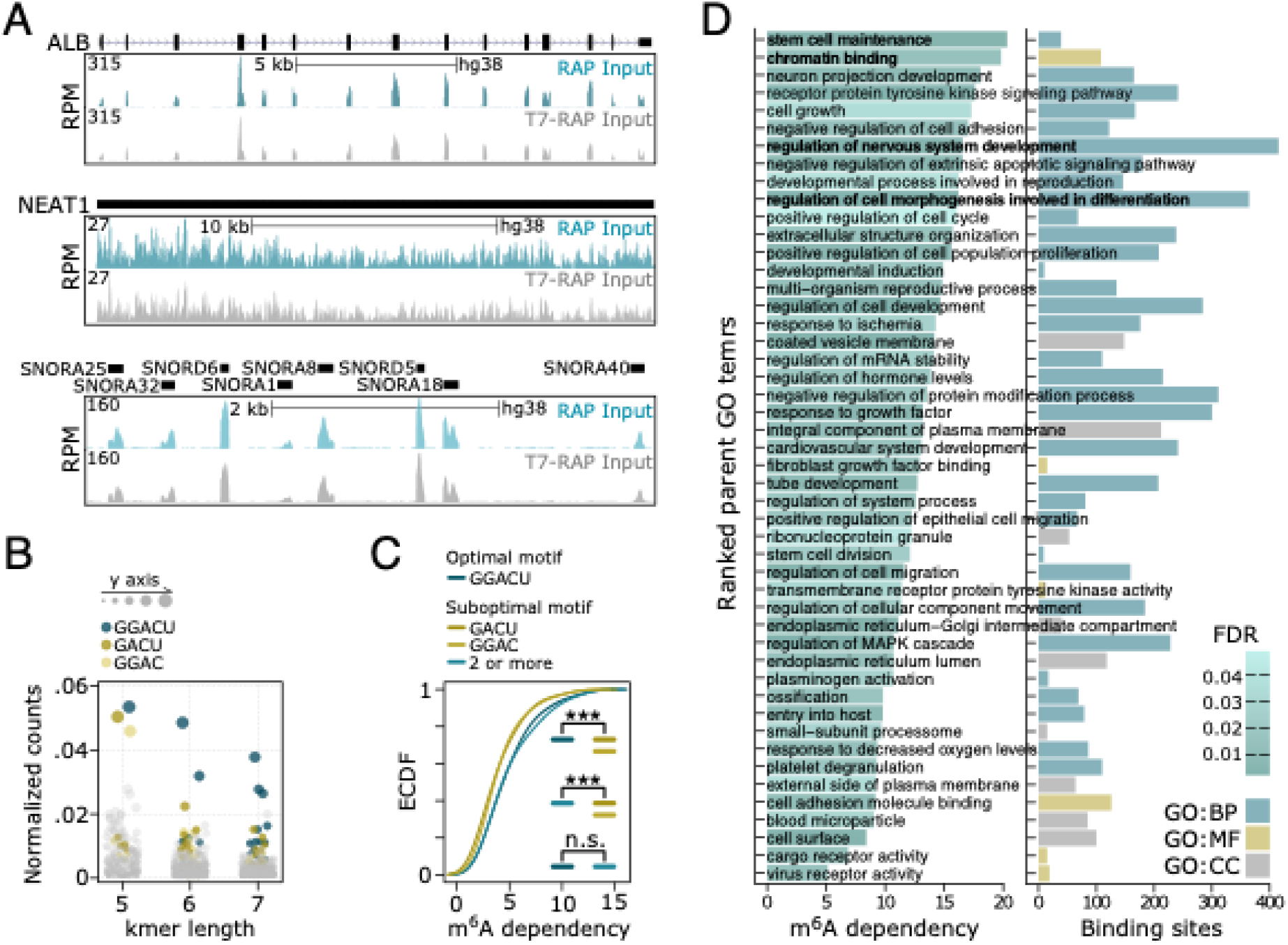
YTHDF1 exhibits extensive m^6^A dependencies as an RNA-binding protein.Related to Figure 5. **(A)** Genome tracks demonstrate similarities between the native (RAP input, blue) and *in vitro* transcribed (T7-RAP input, grey) substrates used for profiling YTHDF1 binding. Tracks exemplify an even transcriptome and fragment representation between the two substrates over a long spliced (top), long unspliced (middle) and small (bottom) transcripts. Genomic locations with scale bar indicate the length of the genomic region in bases and gene features (black rectangle: exon and UTR, grey line: intron, arrow: direction of transcription) (x-axis) and normalized read density (RPM, y-axis) for RAP- and T7-RAPseq are shown. **(B)** Dotplot highlights the enrichment of k-mers of various lengths containing the GGACU motif or its two encompassed 4-mers, GGAC and GACU respectively. The fraction of k-mer counts is weighted the ratio between their abundance at bound sites with respect to control sites (y-axis). **(C)** Line plot of the empirical cumulative distribution functions (ECDF) of YTHDF1 dependency on m^6^A presence in each motif at each binding site. The m^6^A dependency is computed as the ratio between peak binding scores observed at YTHDF1 binding sites when using native RNA or *in vitro* transcribed RNA as substrates. The values are log_2_ transformed. Asterisks represent *p*-values (one-tailed Wilcoxon rank sum test, ***p<0.001). ‘2 or more’ refers to binding of a minimum of two tetramers. **(D)** Bar plots show significantly enriched parent Gene Ontology (GO) terms for YTHDF1-bound genes over the expressed transcriptome (FDR-adjusted *p*≤0.05). GO terms were ranked by m^6^A dependency (left) and the relative number of binding sites that were mapped to each GO term (right). The m^6^A dependency per GO term is represented by the ratio of cumulative peak binding scores per pathway between RAP and T7-RAPseq assays.

**Figure S6.**
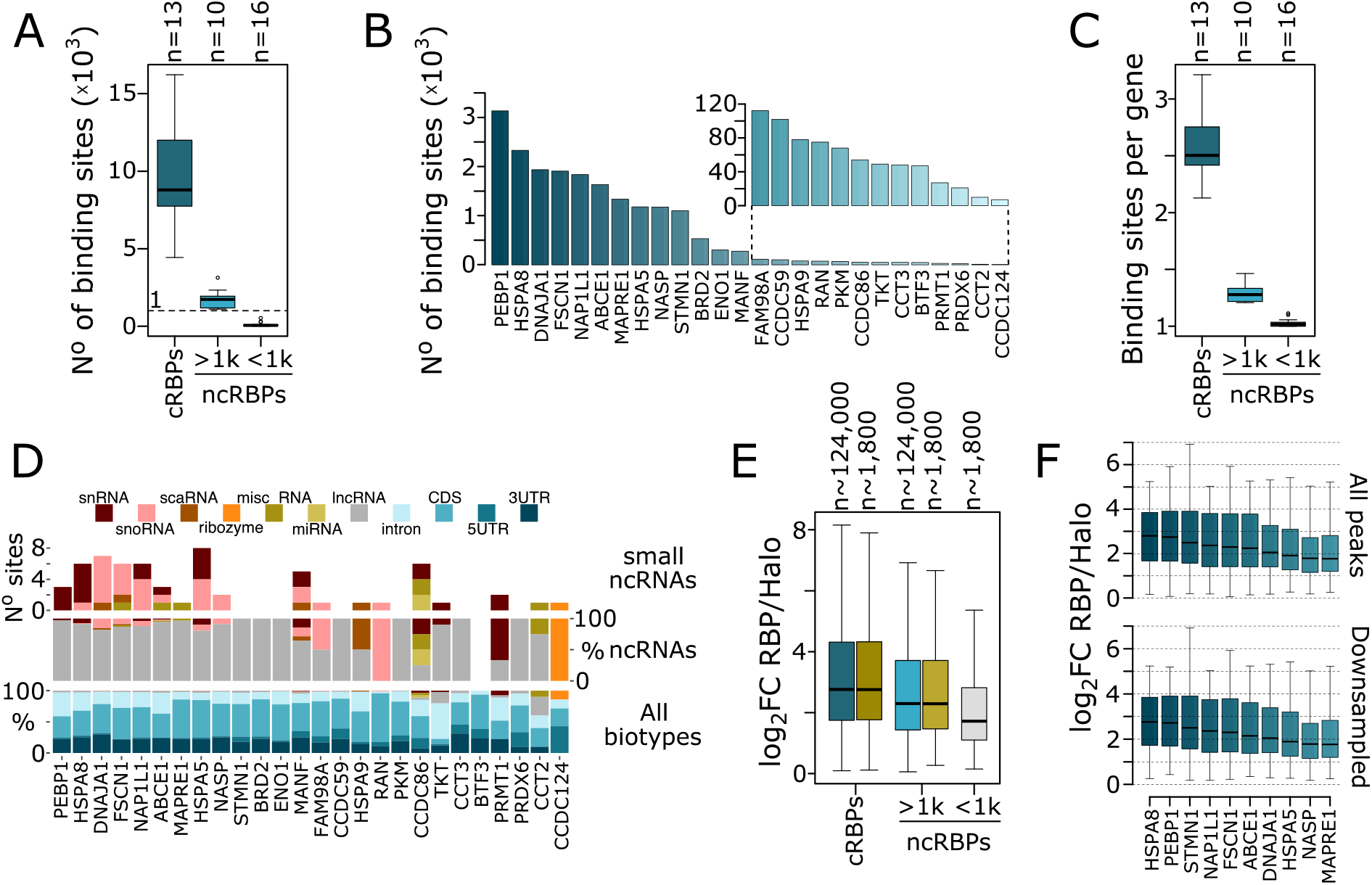
ncRBP have reduced number of endogenous RNA targets compared to cRBPs. Related to Figure 6. **(A-B)** Boxplots show number of (**A**) all binding sites and (**B**) binding sites per gene identified in RAPseq for canonical (cRBPs, dark blue) and noncanonical RBPs (ncRBPs). ncRBPs are split into two groups based on more (light blue) or fewer (grey) than 1000 binding sites. The number (n) of RBPs for each group is indicated. **(C)** Barplots display the number of binding sites identified in RAPseq for all 26 ncRBP. **(D)** Barplots demonstrate the binding site occurrences across RNA biotypes for each ncRBPs mapping to small noncoding RNAs (top, excluding long noncoding RNA), all noncoding RNAs (middle) or all biotypes (bottom). **(E)** Boxplots show the fold-change of enrichments of RBP binding sites (n) over HaloTag control for the three groups. Yellow boxes represent downsampling per group. **(F)** Boxplots display the fold-change of enrichments for ncRBPs with more than 1000 binding sites considering all (top) or downsampled (n=1000) binding sites.

**Figure S7.**
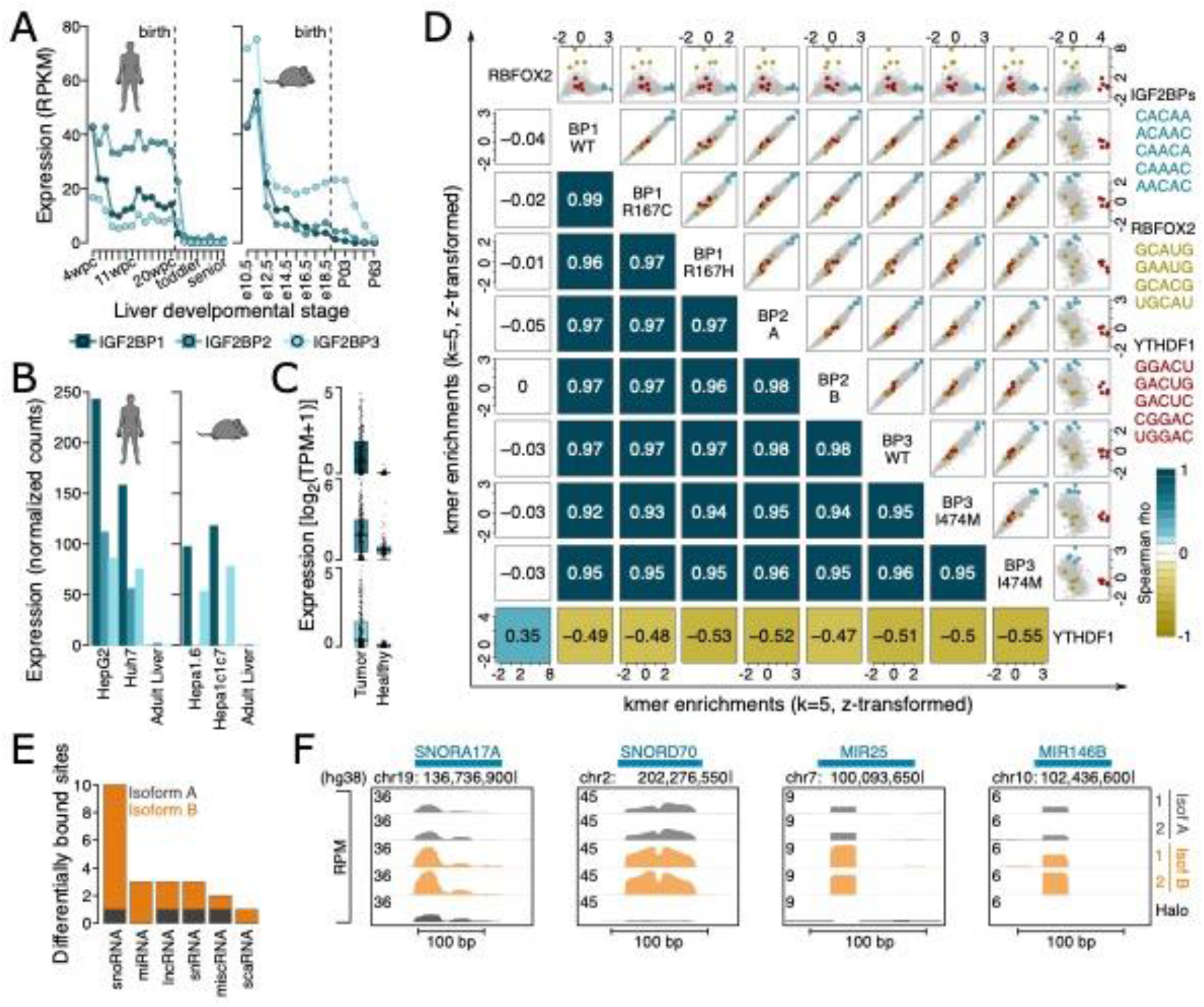
The binding specificity of IGF2BP family disease variants is preserved. Related to Figure 7. **(A)** Line plots show IGF2BP paralog mRNA expression levels (in reads per kilobase per million mapped reads, RPKM) over human (left) and mouse (right) liver development. Birth is indicated by dashed lines. **(B)** Barplots display IGF2BP gene family expression levels in liver cancer cell lines and healthy adult liver in human (left, HepG2 and Huh7) and mouse (right, Hepa1.6 and Hepa1c1c7). **(C)** Boxplots show IGF2BP gene family expression levels in TCGA liver cancer cohorts. Each dot represents one patient. **(D)** Scatter plots (upper-right triangle) illustrate k-mer enrichment correlations between the binding specificities of the three IGF2BP paralogs and their respective variants. RBFOX2 and YTHDF1 are included as controls. Colored dots highlight the binding motifs of each RBP. Spearman rank correlations (ρ) for each comparison are reported (lower-left triangle). Axes present z-transformed k-mer enrichment values. **(E)** Barplot shows the number (y-axis) and type (x-axis) of differentially bound ncRNAs by the two IGF2BP2 isoforms. **(F)** Genome tracks demonstrate IGF2BP2 isoform A (grey) or B (orange) binding over HaloTag control (grey) to two snoRNAs and two miRNAs. Genomic locations with scale bar indicate the length of the genomic region in bases (x-axis) and normalized read density (RPM, y-axis) are shown.

